# Potent Neutralizing Antibodies Directed to Multiple Epitopes on SARS-CoV-2 Spike

**DOI:** 10.1101/2020.06.17.153486

**Authors:** Lihong Liu, Pengfei Wang, Manoj S. Nair, Jian Yu, Micah Rapp, Qian Wang, Yang Luo, Jasper F-W. Chan, Vincent Sahi, Amir Figueroa, Xinzheng V. Guo, Gabriele Cerutti, Jude Bimela, Jason Gorman, Tongqing Zhou, Zhiwei Chen, Kwok-Yung Yuen, Peter D. Kwong, Joseph G. Sodroski, Michael T. Yin, Zizhang Sheng, Yaoxing Huang, Lawrence Shapiro, David D. Ho

**Affiliations:** Aaron Diamond AIDS Research Center, Columbia University Vagelos College of Physicians and Surgeons, New York, NY 10032, USA; Zuckerman Mind Brain Behavior Institute, Columbia University, New York, NY 10027, USA; Dana-Farber Cancer Institute, Harvard Medical School, Boston, MA 02215, USA; State Key Laboratory of Emerging Infectious Diseases, Carol Yu Centre for Infection, Department of Microbiology, Li Ka Shing Faculty of Medicine, The University of Hong Kong, Hong Kong Special Administrative Region, China; Centre for Virology, Vaccinology and Therapeutics, Health@InnoHK, The University of Hong Kong, Hong Kong Special Administrative Region, China; Department of Microbiology & Immunology Flow Cytometry Core, Columbia University Vagelos College of Physicians and Surgeons, New York, NY 10032, USA; Human Immune Monitoring Core, Columbia University Vagelos College of Physicians and Surgeons, New York, NY 10032, USA; Vaccine Research Center, National Institutes of Health, Bethesda, MD 20892, USA; AIDS Institute, Li Ka Shing Faculty of Medicine, The University of Hong Kong, Hong Kong Special Administrative Region, China; Department of Biochemistry, Columbia University, New York, NY 10032, USA; Division of Infectious Diseases, Department of Internal Medicine, Columbia University Vagelos College of Physicians and Surgeons, New York, NY 10032, USA

## Abstract

The SARS-CoV-2 pandemic rages on with devasting consequences on human lives and the global economy^1,2^. The discovery and development of virus-neutralizing monoclonal antibodies could be one approach to treat or prevent infection by this novel coronavirus. Here we report the isolation of 61 SARS-CoV-2-neutralizing monoclonal antibodies from 5 infected patients hospitalized with severe disease. Among these are 19 antibodies that potently neutralized the authentic SARS-CoV-2 *in vitro*, 9 of which exhibited exquisite potency, with 50% virus-inhibitory concentrations of 0.7 to 9 ng/mL. Epitope mapping showed this collection of 19 antibodies to be about equally divided between those directed to the receptor-binding domain (RBD) and those to the N-terminal domain (NTD), indicating that both of these regions at the top of the viral spike are immunogenic. In addition, two other powerful neutralizing antibodies recognized quaternary epitopes that are overlapping with the domains at the top of the spike. Cryo-electron microscopy reconstructions of one antibody targeting RBD, a second targeting NTD, and a third bridging two separate RBDs revealed recognition of the closed, “all RBD-down” conformation of the spike. Several of these monoclonal antibodies are promising candidates for clinical development as potential therapeutic and/or prophylactic agents against SARS-CoV-2.

## Background

A novel coronavirus, now termed SARS-CoV-2^1,2^, has caused over 11 million confirmed infections globally, leading to over 500,000 deaths. This pandemic has also put much of the world on pause, with unprecedented disruption of lives and unparalleled damage to the economy. A return to some semblance of normalcy will depend on science to deliver an effective solution, and the scientific community has responded admirably. Drug development is well underway, and vaccine candidates have entered clinical trials. Another promising approach is the isolation of SARS-CoV-2-neutralizing monoclonal antibodies (mAbs) that could be used as therapeutic or prophylactic agents. The primary target for such antibodies is the viral spike, a trimeric protein^3,4^ that is responsible for binding to the ACE2 receptor on the host cell^1,3,5,6^. The spike protein is comprised of two subunits. The S1 subunit has two major structural elements: RBD and NTD; the S2 subunit mediates virus-cell membrane fusion after the RBD engages ACE2. Reports of discovery of neutralizing mAbs that target the RBD have been published recently^7-11^. We now describe our efforts in isolating and characterizing a collection of mAbs that not only target multiple epitopes on the viral spike but also show exquisite potency in neutralizing SARS-CoV-2.

### Patient Selection

Forty patients with PCR-confirmed SARS-CoV-2 infection were enrolled in a cohort study on virus-neutralizing antibodies. Plasma samples from all subjects were first tested for neutralizing activity against SARS-CoV-2 pseudovirus (Wuhan-Hu-1 spike pseudotyped with vesicular stomatitis virus). Widely varying neutralizing titers were observed, with IC_50_ ranging from a reciprocal plasma dilution of <100 to ∼13,000 (Fig. 1a). Five patients were chosen for mAb isolation because their plasma virus-neutralizing titers were among the highest. The clinical characteristics of these 5 cases are summarized in Extended Data Table 1. All were severely ill with acute respiratory distress syndrome requiring mechanical ventilation.

**Fig. 1.**
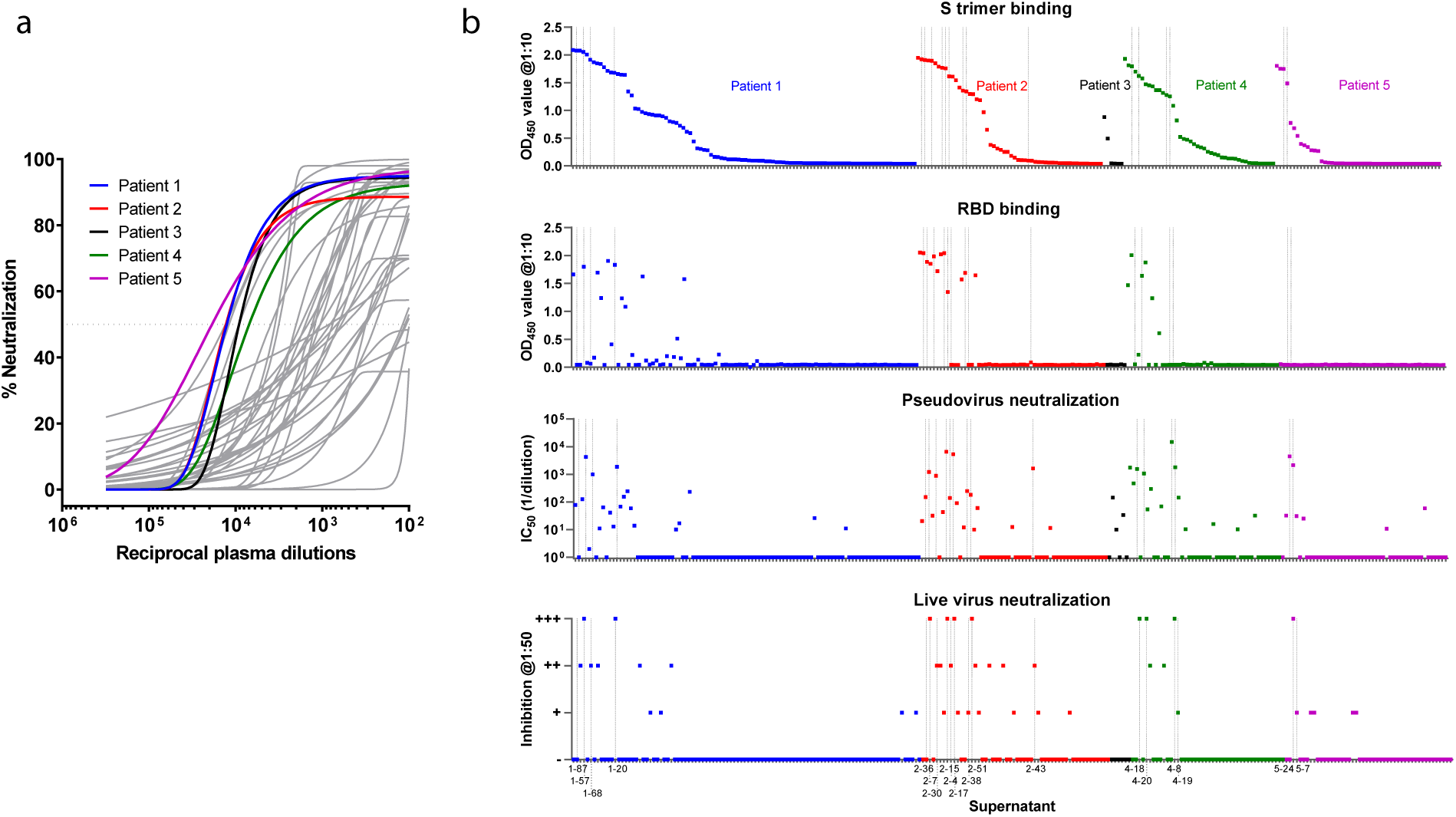
Isolation of SARS-CoV-2 mAbs from infected patients with severe disease. **a**, Plasma neutralization profile of 40 patients against SARS-CoV-2 pseudovirus (highlighted are 5 top neutralizers chosen). **b**, All 252 transfection supernatants were screened for binding to S trimer and RBD, as well as for neutralization against SARS-CoV-2 pseudovirus and live virus. For pseudovirus neutralization, the 50% inhibitory dilutions (IC_50_) of each supernatant were plotted. For live virus, semi-quantitative representation of the inhibition at a dilution of 1:50, with neutralization levels ranging from (-) for none to (+++) for complete neutralization, was plotted. Potent antibodies later identified are marked by vertical lines and labelled at the bottom. The antibodies from each patient are colored as in **a**.

### Monoclonal Antibody Isolation and Construction

Peripheral blood mononuclear cells from each patient were put through an experimental schema (Extended Data Fig. 1a) starting with cell sorting by flow cytometry. The sorting strategy focused on live memory B lymphocytes that were CD3-negative, CD19-positive, and CD27-positive (Extended Data Fig. 1b). The final step focused on those cells that bound the SARS-CoV-2 spike trimer (S trimer)^4^. A total of 602, 325, 14, 147, and 145 such B cells from Patient 1, Patient 2, Patient 3, Patient 4, and Patient 5, respectively, were labelled with a unique hashtag (Extended Data Fig. 1c). The cells were then placed into the 10X Chromium (10X Genomics) for single-cell 5’mRNA and V(D)J sequencing to obtain paired heavy (H) and light (L) chain sequences. A careful bioinformatic analysis was carried out on 1,145 paired sequences to downselect “high-confidence” antigen-specific mAbs. A total of 331 mAb sequences were recovered, representing 252 individual clones. Only 6 mAbs were from Patient 3, whereas 44 to 100 mAbs were identified from each of the other patients (Extended Data Table 2). The VH and VL sequences of 252 antibodies (one per clone) were codon-optimized and synthesized, and each VH and VL gene was then cloned into an expression plasmid with corresponding constant region of H chain and L chain of human IgG1, and mAbs were expressed by co-transfection of paired full-length H chain and L chain genes into Expi293 cells.

### Monoclonal Antibody Screening

All 252 transfection supernatants were screened for binding to S trimer and RBD by enzyme-linked immunosorbent assays (ELISAs), as well as for neutralization against SARS-CoV-2 pseudovirus and live virus. These results are graphically represented in Fig. 1b and tabulated in Extended Data Table 2. It was evident that a substantial percentage of the mAbs in the supernatants bound S trimer, and a subset of those bound RBD. Specifically, 121 supernatants were scored as positive for S trimer binding, yielding an overall hit rate of 48%. Of these, 38 were positive for RBD binding while the remaining 83 were negative. None of the 13 trimer-specific mAbs from Patient 5 recognized RBD. In the pseudovirus neutralization screen, 61 supernatants were scored as positive, indicating that half of the trimer-specific mAbs were virus neutralizing. In the screen for neutralization against SARS-CoV-2 (strain USA-WA1/2020), 41 supernatants were scored as positive. Overall, this screening strategy was quite effective in picking up neutralizing mAbs (vertical lines and labelled antibodies at the bottom of Fig. 1b) that were later identified as potent.

### Sequence Analysis of S Trimer-Specific Monoclonal Antibodies

Of the 121 mAbs that bound S trimer, 88% were IgG isotype, with IgG1 being predominant (Extended Data Fig. 2a). Comparison to the IgG repertoire of three healthy human donors^12^ revealed a statistically significant over-representation of IGHV3-30, IGKV3-20, and IGHJ6 genes for this collection of SARS-CoV-2 mAbs (Extended Data Figs. 2b and 2c). A longer average CDRH3 length was also noted (Extended Data Fig. 2d). Interestingly, the average percentages of somatic hypermutation in VH and VL were 2.1 and 2.5, respectively, which were significantly lower than those found in healthy individuals (Extended Data Fig. 2e) and remarkably close to germline sequences.

### Antigen Binding and Virus Neutralization

Since the screening for pseudovirus neutralization was performed quantitatively with serial dilutions of the transfection supernatants, we plotted in Extended Data Fig. 3 the best-fit neutralization curves for 130 samples that were positive in at least one of the screens shown in Fig. 1b. Most were non-neutralizing or weakly neutralizing, but 18 showed evidently better potency.

One additional supernatant was initially missed in the pseudovirus screen (Patient 1 in Extended Data Fig. 3) but was later found to be a potent neutralizing mAb. Together, these 19 mAbs were purified from transfection supernatants and further characterized for their binding and neutralization properties. As shown in Fig. 2a, all but one (2-43) of the mAbs bound the S trimer by a quantitative ELISA. Nine of the antibodies clearly bound RBD, with little or no binding to NTD. Eight antibodies bound NTD to varying degrees, with no binding to RBD. Two mAbs bound neither RBD nor NTD, and were therefore categorized as “Others”.

**Fig. 2.**
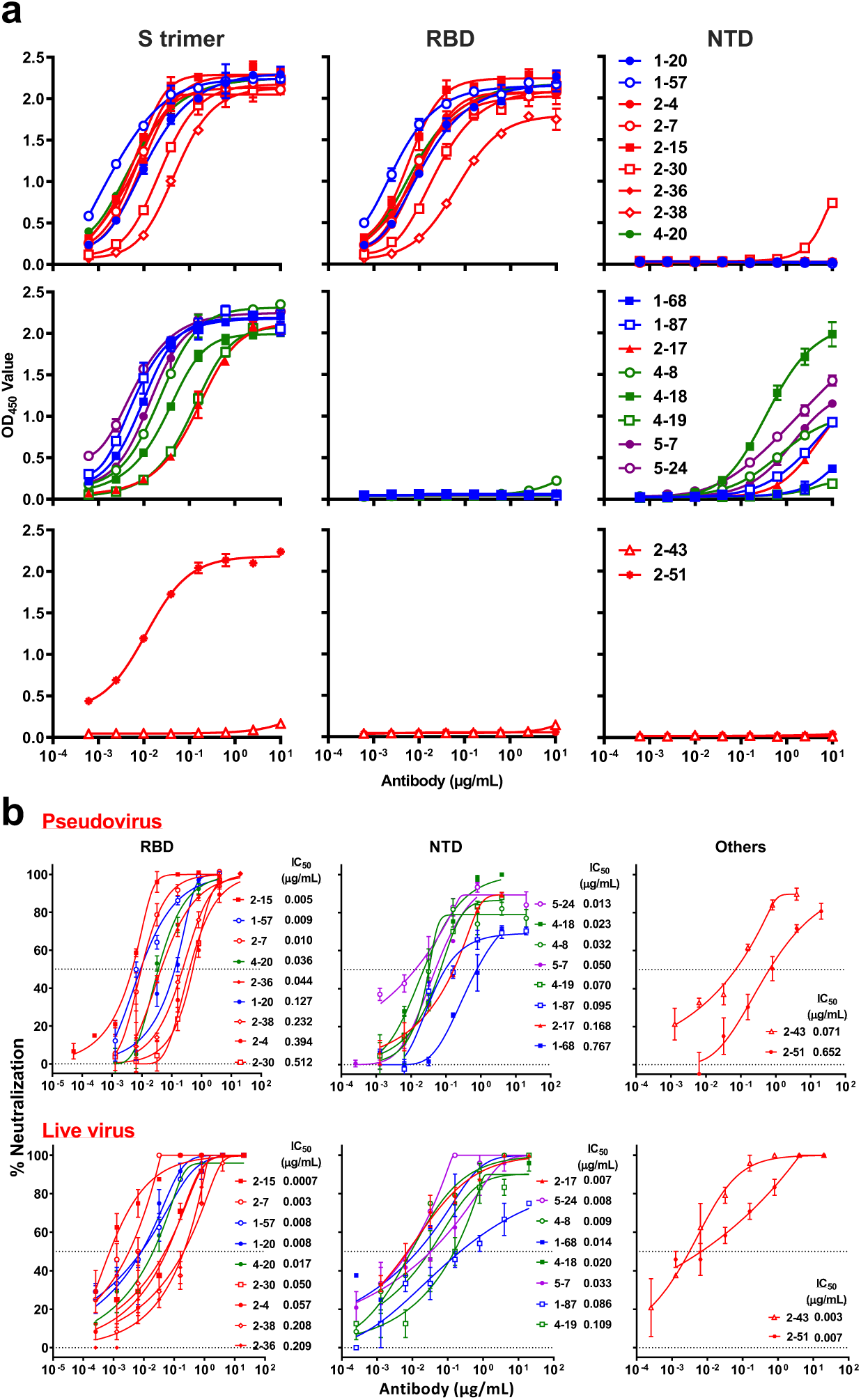
Characterization of SARS-CoV-2 potent neutralizing mAbs. **a**, Binding profiles of 19 purified potent neutralizing mAbs against SARS-CoV-2 S trimer (left), RBD (middle), and NTD (right). Note that mAb 2-30 bound multiple proteins at high concentrations. **b**, The pseudovirus (top panels) and live virus (bottom panels) neutralization profiles for the 19 purified mAbs. Epitope classifications are listed on top of the panel b. Single replicate of the binding experiment and triplicates of neutralization are presented as mean ± SEM.

The pseudovirus neutralization profiles for these purified 19 mAbs are shown in the top portion of Fig. 2b. The RBD-directed antibodies neutralized the pseudovirus with IC_50_ of 0.005 to 0.512 μg/mL; the NTD-directed antibodies were slightly less potent, with IC_50_ ranging from 0.013 to 0.767 μg/mL. A common feature of the NTD mAbs was the plateauing of virus neutralization at levels short of 100%. Two antibodies, categorized as “Others”, neutralized with IC_50_ of 0.071 and 0.652 μg/mL. Antibody neutralization of the authentic or live SARS-CoV-2 (strain USA-WA1/2020) was carried out using Vero cells inoculated with a multiplicity of infection of 0.1. As shown in the bottom portion of Fig. 2b, the RBD-directed antibodies again neutralized the virus but with IC_50_ of 0.0007 to 0.209 μg/mL; the NTD-directed antibodies showed similar potency, with IC_50_ ranging from 0.007 to 0.109 μg/mL. Here, the plateauing effect seen in the pseudovirus neutralization assay was less apparent. Antibodies 2-43 and 2-51 neutralized the live virus with IC_50_ of 0.003 and 0.007 μg/mL, respectively. Overall, nine mAbs exhibited exquisite potency in neutralizing authentic SARS-CoV-2 *in vitro* with IC_50_ of 0.009 μg/mL or less, including four directed to RBD (2-15, 2-7, 1-57, and 1-20), three to NTD (2-17, 5-24, and 4-8), and two to undetermined regions on the S trimer (2-43 and 2-51). It is remarkable that Patient 2 alone contributed five of the top nine SARS-CoV-2 neutralizing mAbs. Correlation of the results of the two virus-neutralizing assays is shown in Extended Data Fig. 4.

### Epitope Mapping

All 19 potent neutralizing mAbs (Fig. 2) were further studied in antibody competition experiments to gain insight into their epitopes. In addition, 12 mAbs that bound the S trimer strongly during the initial supernatant screen were also chosen, including 5 that bound RBD (1-97, 2-26, 4-13, 4-24, and 4-29) and 7 that did not bind RBD (1-21, 2-29, 4-15, 4-32, 4-33, 4-41, and 5-45). Four of these mAbs were weak in neutralizing SARS-CoV-2 pseudovirus, and the remaining 8 were non-neutralizing (Extended Data Fig. 5). First, a total of 16 non-RBD mAbs were evaluated for competition in binding to S trimer by ELISA in a “checkerboard” experiment. The extent of the antibody competition is reflected by the intensity of the heatmap shown in Fig. 3a. There is one large cluster (A) of mAbs that competed with one another, and it partially overlaps a small cluster (B). A third cluster (C) does not overlap at all. Note that all but one of the antibodies in cluster A recognized NTD. Antibody 2-51 is clearly directed to the NTD region even though it could not bind NTD. Moreover, one mAb each from clusters B and C also recognized NTD, thereby indicating that all three clusters are within or near the NTD. One mAb, 1-21, appears to have a unique non-overlapping epitope (epitope region D).

**Fig. 3.**
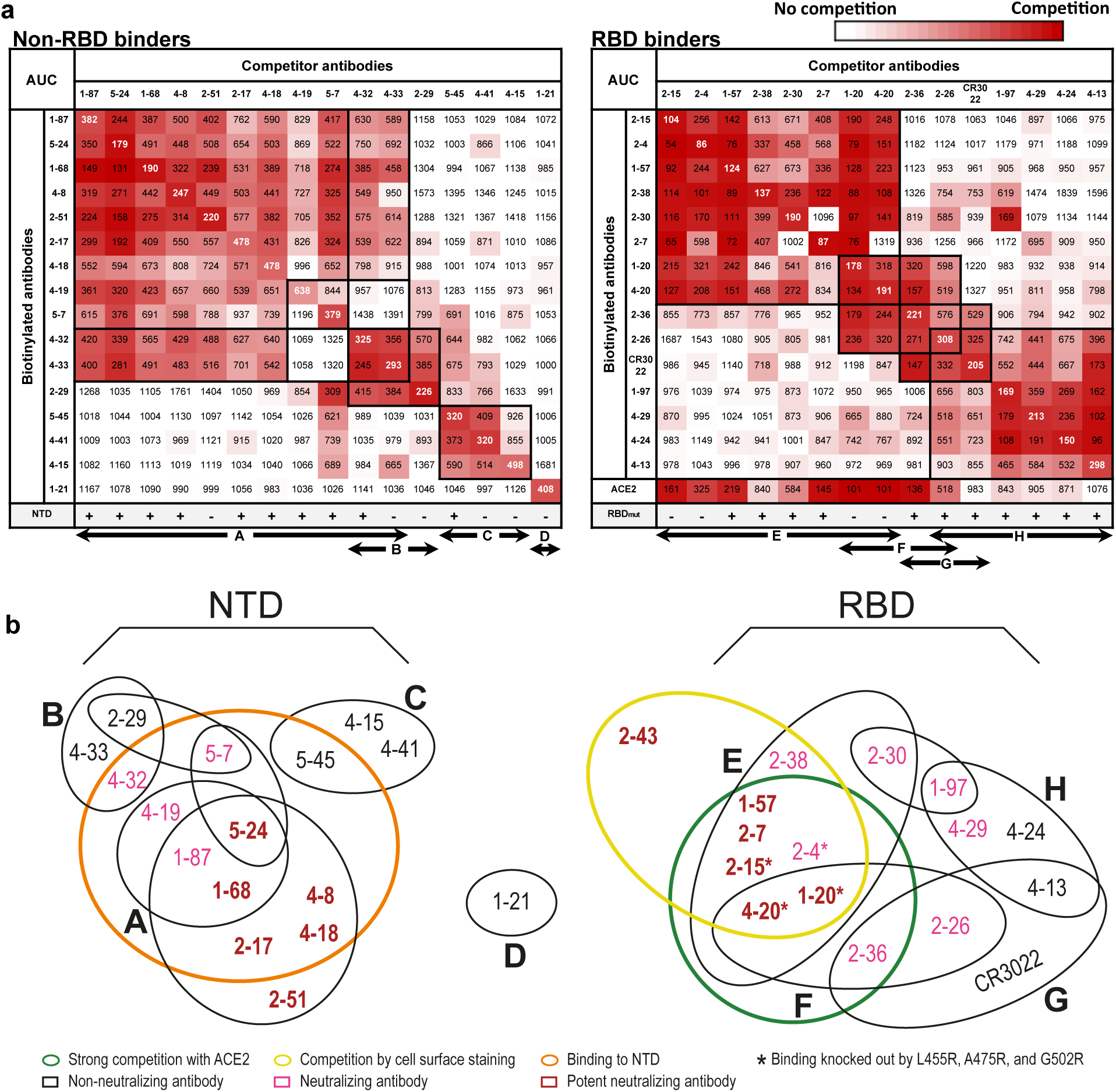
Epitope mapping of select neutralizing and non-neutralizing mAbs. **a**, competition results of non-RBD binders (left) and RBD binders (right) in blocking ACE2 or biotinylated mAb binding to the S trimer. In addition, the ability to bind NTD and RBD_mut_ of each mAb is shown. The numbers in each box show the area under each competition curve (AUC) as tested by ELISA. +/- indicates binding/no binding of the mAb to the protein. The letters A to H at the bottom denote clusters of antibody epitopes defined by the competition experiments. **b**, Venn diagram interpretation of results from **a** and Extended Data Fig. 6b.

Second, a similar “checkerboard” competition experiment was carried out by ELISA for 14 of our RBD-directed mAbs plus CR3022^13,14^. Here, the heatmap shows that there are four epitope clusters that are serially overlapping (Fig. 3a). There is one large cluster (E) that contains mAbs largely capable of blocking ACE2 binding. Furthermore, 4 antibodies in this cluster lost binding to a mutant RBD (L455R, A475R, G502R) that could no longer bind ACE2 (our unpublished findings). Taken together, these results suggest that most of the mAbs in cluster E are directed to the ACE2-binding interface of RBD. The next cluster (F) connects to both cluster E and cluster G, the location of which is defined by its member CR3022^15^. Lastly, cluster G overlaps another cluster (H), which includes 1-97 that strongly inhibited the binding of 2-30 to the S trimer. This finding suggests that cluster H may be proximal to one edge of cluster E.

One potent neutralizing mAb, 2-43, did not bind S trimer by ELISA (Fig. 2a) and thus could not be tested in the above competition experiments. However, 2-43 did strongly bind S trimer when expressed on the cell surface as determined by flow cytometry (Extended Data Fig. 6a), and this binding was competed out by itself but not by RBD, NTD, ACE2, or the soluble S trimer^4^ (Extended Data Fig. 6b). NTD-directed mAbs had only a modest effect on its binding to cell-surface S trimer, but numerous RBD-directed mAbs in cluster E potently blocked the binding of 2-43, demonstrating that this antibody is likely targeting a quaternary epitope on the top of RBD.

These mapping results could be represented by two sets of Venn diagrams shown in Fig. 3b. In the non-RBD region, the potent neutralizing mAbs reside exclusively in cluster A and bind a patch on the NTD. Weaker neutralizing mAbs recognize a region at the interface between clusters A and B. In the RBD region, the most potent neutralizing mAbs also group together within one cluster (E). Given that all block ACE2 binding, it is likely they recognize the top of RBD and neutralize the virus by competitive inhibition of receptor binding. Cluster G contains CR3022, a mAb known to be directed to an epitope on a cryptic site on the side of RBD when it is in the “up” position^15^. Cluster F is therefore likely situated between the top and this “cryptic” site. The Venn diagram also suggests that cluster H may occupy a different side surface of RBD, perhaps in the region recognized by S309, a mAb isolated from a SARS-CoV-1 patient^8^.

### Cryo-Electron Microscopy

We produced cryo-EM reconstructions of Fabs from three mAbs in complex with the S trimer^4^. First, single-particle analysis of the complex with 2-4 Fab (RBD-directed) yielded maps of high quality (Fig. 4a; Extended Data Table 3; Extended Data Fig. 7), with the most abundant particle class representing a 3-Fab-per-trimer complex, refined to an overall resolution of 3.2 Å. While density for the constant portion of the Fabs was visible, it was blurred due to molecular motion, and thus only the variable domains were included in the molecular model. Fab 2-4 bound the spike protein near the apex, with all RBDs in the “down” orientation, and the structure of the antibody-bound spike protein was highly similar to previously published unliganded spike structures in the “all-down” conformation^3,4^. Detailed interactions between 2-4 and RBD are discussed and exhibited in Extended Data Fig. 8. Overall, the structure of the 2-4 Fab-spike complex shows that neutralization of SARS-CoV-2 by this mAb likely results from locking RBD in the down conformation while also occluding access to ACE2.

**Figure 4.**
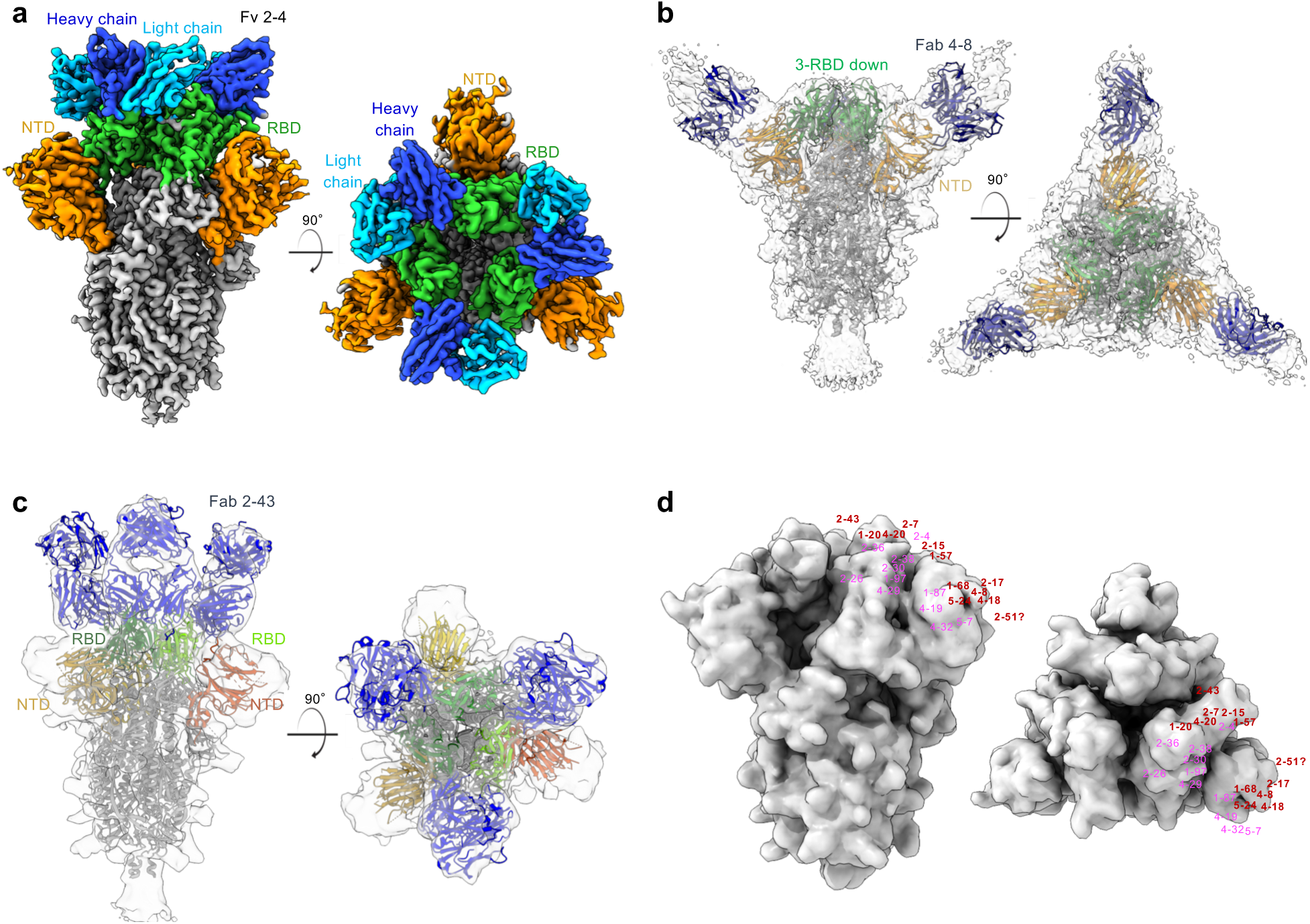
Cryo-EM reconstructions of Fab-spike complexes and visualization of neutralizing epitopes on the spike surface. **a**, Cryo-EM reconstruction of 2-4 Fab in complex with S trimer at 3.2 Å overall resolution. Density is colored with RBD in green, NTD in orange, and other regions in grey. **b**, Cryo-EM reconstruction of 4-8 Fab in complex with S trimer (ribbon diagram, colored as in a) at 3.9 Å overall resolution, with RBDs in the “all-down” configuration. **c**, Cryo-EM reconstruction of the 2-43 Fab in complex with S trimer at 5.8 Å resolution reveals a quaternary epitope involving RBD from one subunit and another RBD from the next. **d**, Mapping of the Venn diagrams from Fig. 3b onto the surface of the viral spike.

Second, we also produced 3D cryo-EM reconstructions of 4-8 Fab (NTD-directed) in complex with the S trimer (Extended Data Table 3; Extended Data Fig. 9). Two main particle classes were observed – one for a 3-Fab-bound complex with all RBDs “down” at 3.9 Å resolution (Fig. 4b), and another a 3-Fab-bound complex with one RBD “up” at 4.0 Å resolution (Extended Data Fig. 10). However, molecular motion prevented visualization of the interaction at high resolution. Nevertheless, the density in the 4-8 map reveals the overall positions of the antibody chains targeting NTD. Such binding to the tip of NTD results in SARS-CoV-2 neutralization remains unclear.

Third, a 5.8 Å resolution reconstructions of 2-43 Fab in complex with the S trimer (Extended Data Table 3; Extended Data Fig. 11) revealed three bound Fabs, each targeting a quaternary epitope on the top of the spike that included elements of the RBDs from two adjacent S1 protomers (Fig. 4c), consistent with the epitope mapping results (Extended Data Fig. 6b and Fig. 3b), including the lack of binding to isolated RBD (Fig. 2a). Given these findings, the inability of 2-43 to bind S trimer by ELISA is likely an artifact of the assay format, as this mAb did bind the spike expressed on the cell surface and in the cryo-EM study.

Armed with these three cryo-EM reconstructions, we used the Venn diagrams from Fig. 3b to map the epitopes of many of our SARS-CoV-2 neutralizing mAbs onto the surface of the spike (Fig. 4d). This is obviously a rough approximation since antibody footprints are much larger than the area occupied by the mAb number. However, the spatial relationship of the antibody epitopes should be reasonably represented by Fig. 4d.

### Protection of Hamsters from SARS-CoV-2 Infection by mAb 2-15

To assess the *in vivo* potency of mAb 2-15, we performed a protection experiment in a golden Syrian hamster model of SARS-CoV-2 infection. The hamsters were first given an intraperitoneal injection of the antibody at a dose of 1.5 mg/kg or 0.3 mg/kg, or PBS alone. Intranasal inoculations of 10^5^ plaque-forming units (PFU) of the HKU-001a strain of SARS-CoV-2 were carried out 24 hours later. Four days after virus challenge, lung tissues were harvested to quantify the viral load.

As shown in Fig. 5, both viral RNA copy numbers and infectious virus titers were reduced by 4 logs or more in hamsters given the dose of 1.5 mg/kg of mAb 2-15. The protection at the dose of 0.3 mg/kg was borderline, as we had estimated. This pilot animal study demonstrates that the potency of mAb 2-15 *in vitro* is indeed reflected *in vivo*, with complete elimination of infectious SARS-CoV-2 at a relatively modest antibody dose.

**Figure 5.**
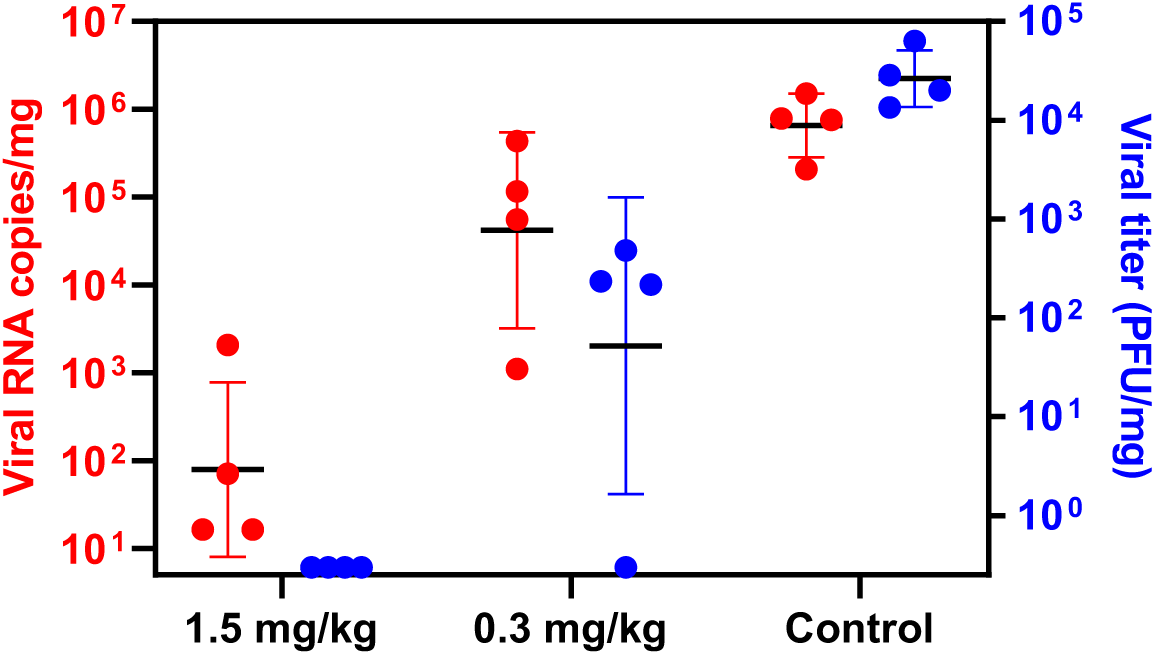
Efficacy of mAb 2-15 in protecting against SARS-CoV-2 infection in lung tissues of hamsters. One day prior to intranasal challenge of SARS-CoV-2, each group of hamsters was given a single intraperitoneal dose of 1.5 mg/kg of mAb 2-15 (n = 4), 0.3 mg/kg of mAb 2-15 (n = 4), or saline as control (n = 4). The viral loads in the lung tissues on day 4 after viral challenge were determined by qRT-PCR (red), as well as by an assay to quantify plaque forming units (PFU) of infectious SARS-CoV-2 (blue). All data points are shown, along with the mean ± SD. The differences between 1.5 mg/kg group and controls are statistically significant with p value <0.05.

## Discussion

We have discovered a collection of SARS-CoV-2-neutralizing mAbs that are not only potent but also diverse. Nine of these antibodies can neutralize the authentic virus *in vitro* at concentrations of 9 ng/mL or less (Fig. 2b), including 4 directed to RBD, 3 directed to NTD, and 2 to quaternary epitopes nearby. Surprisingly, many of the these mAbs have V(D)J sequences close to germline sequences, without extensive somatic hypermutations (Extended Data Fig. 2e), a finding that bodes well for vaccine development. Our most potent RBD-specific mAbs (e.g., 2-15, 2-7, 1-57, and 1-20) compare favorably with such antibodies recently reported^7,8,10,16-20^, including those with high potency^9,11,21,22^. The *in vitro* potency of 2-15 is well reflected *in vivo* in the hamster protection experiment (Fig. 5). It appears from the epitope mapping studies that mAbs directed to the top of RBD strongly compete with ACE2 binding and potently neutralize the virus, whereas those directed to the side surfaces of RBD do not compete with ACE2 and neutralize less potently (Figs. 3b and 4d). Our collection of non-RBD neutralizing mAbs is unprecedented in that such antibodies only have been sporadically reported and only with substantially lower potencies^22-24^. The most potent of these mAbs are directed to (e.g., 2-17, 5-24, and 4-8) or overlapping with (2-51) a patch on the NTD (Figs. 3b and 4d). It is unclear how NTD-directed mAbs block SARS-CoV-2 infection and why their neutralization profiles are different from those of RBD-directed antibodies (Fig. 2b). Nevertheless, vaccine strategies that do not include NTD will be unable to induce an important class of virus-neutralizing antibodies.

The isolation of two mAbs (2-43 and 2-51) directed to epitopes that do not map to RBD and NTD is also unprecedented. Cryo-EM of 2-43 Fab bound to the S trimer has confirmed its epitope as quaternary in nature, crossing from the top of one RBD to the top of another RBD (Fig. 4c). It will be equally informative to understand the epitope of 2-51. In this study, we also show the first evidence by cryo-EM for a neutralizing mAb (4-8) bound to the NTD of the viral spike (Fig. 4b), as well as another high-resolution structure of a mAb (2-4) bound to RBD (Fig. 4a).

The potency and diversity of our SARS-CoV-2-neutralizing mAbs are likely attributable to patient selection. Previously, we observed that infected individuals with severe disease developed a more robust virus-neutralizing antibody response^25^. If Patient 2 had not been included, five of the top neutralizing mAbs would be lost. The diversity of our antibodies is also attributable, in part, to the choice of using the S trimer to sort from memory B cells, while most groups focused on the use of RBD^7,9-11,16-19,21^. The characterization of this diverse collection of mAbs has allowed us to observe that all potent SARS-CoV-2-neutralizing antibodies described to date are directed to the top of the viral spike. RBD and NTD are, no doubt, quite immunogenic. Neutralizing antibodies to the stem region of the S trimer remain to be discovered. In conclusion, we believe several of our monoclonal antibodies with exquisite virus-neutralizing activity are promising candidates for development as modalities to treat or prevent SARS-CoV-2 infection.

## Methods

### Expression and Purification of SARS-CoV-2 Proteins

The mammalian expression vector that encodes the ectodomain of the SARS-CoV-2 S trimer and the vector encoding RBD fused with SD1 at the N-terminus and an HRV-3C protease cleavage site followed by a mFc tag and an 8xHis tag at the C-terminus were kindly provided by Jason McLellan^4^. SARS-CoV-2 NTD (aa1-290) with an HRV-3C protease cleavage site, a mFc tag, and an 8xHis tag at the C-terminus was also cloned into mammalian expression vector pCAGGS. Each expression vector was transiently transfected into Expi293 cells using 1 mg/mL of polyethylenimine (Polysciences). Five days post transfection, the S trimer was purified using Strep-Tactin XT Resin (Zymo Research), and the RBD-mFc and NTD-mFc were purified using protein A agarose (ThermoFisher Scientific). In order to obtain RBD-SD1 and NTD, the mFc and 8xHis tags at the C-terminus were removed by HRV-3C protease (Millipore-Sigma) and then purified using Ni-NTA resin (Invitrogen) followed by protein A agarose.

### Sorting for S Trimer-Specific B cells and Single-Cell BCR Sequencing

Peripheral blood mononuclear cells from five patients and one healthy donor were stained with LIVE/DEAD™ Fixable Yellow Dead Cell Stain Kit (Invitrogen) at ambient temperature for 20 mins, followed by washing with RPMI-1640 complete medium and incubation with 10 µg/mL of S trimer at 4°C for 45 mins. Afterwards, the cells were washed again and incubated with a cocktail of flow cytometry and hashtag antibodies, containing CD3 PE-CF594 (BD Biosciences), CD19 PE-Cy7 (Biolegend), CD20 APC-Cy7 (Biolegend), IgM V450 (BD Biosciences), CD27 PerCP-Cy5.5 (BD Biosciences), anti-His PE (Biolegend), and human Hashtag 3 (Biolegend) at 4°C for 1 hr. Stained cells were then washed, resuspended in RPMI-1640 complete medium and sorted for S trimer-specific memory B cells (CD3-CD19+CD27+S trimer+ live single lymphocytes). The sorted cells were mixed with mononuclear cells from the same donor, labeled with Hashtag 1, and loaded into the 10X Chromium chip of the 5’ Single Cell Immune Profiling Assay (10X Genomics) at the Columbia University Human Immune Monitoring Core (HIMC; RRID:SCR_016740). The library preparation and quality control were performed according to manufacturer’s protocol and sequenced on a NextSeq 500 sequencer (Illumina).

### Identification of S Trimer-Specific Antibody Transcripts

For each sample, full-length antibody transcripts were assembled using the VDJ module in Cell Ranger (version 3.1.0, 10X Genomics) with default parameters and the GRCh38 genome as reference. To identify cells from the antigen sort, we first used the count module in Cell Ranger to calculate copies of all hashtags in each cell from the Illumina NGS raw reads. High confidence antigen-specific cells were identified as follows. Briefly, based on the copy numbers of the hashtags observed, a cell must contain more than 100 copies of the antigen sort-specific hashtag to qualify as an antigen-specific cell. Because hashtags can fall off from cells and bind to cells from a different population in the sample mixture, each cell usually has both sorted and spiked-in-specific hashtags. To enrich for true antigen-specific cells, the copy number of the specific hashtag has to be at least 1.5x higher than that of the non-specific hashtag. Low quality cells were identified and removed using the cell-calling algorithm in Cell Ranger. Cells that do not have productive H and L chain pairs were excluded. If a cell contains more than two H or/and L chain transcripts, the transcripts with less than 3 unique molecular identifiers were removed. Cells with identical H and L chain sequences, which may have resulted from mRNA leakage, were merged into one cell. Additional filters were applied to remove low quality cells and/or transcripts in the antibody gene annotation process.

### Antibody Transcript Annotation and Selection Criteria

Antigen-specific antibody transcripts were processed using our bioinformatics pipeline SONAR for quality control and annotation^26^. Briefly, V(D)J genes were assigned for each transcript using BLAST^27^ with customized parameters against a germline gene database obtained from the international ImMunoGeneTics information system (IMGT) database^26,28^. Based on BLAST alignments of V and J regions, CDR3 was identified using the conserved second cysteine in the V region and WGXG (H chain) or FGXG (L chain) motifs in the J region (X represents any amino acid). For H chain transcripts, the constant domain 1 (CH1) sequences were used to assign isotype using BLAST with default parameters against a database of human CH1 genes obtained from IMGT. A BLAST E-value threshold of 1E-6 was used to find significant isotype assignments, and the CH1 allele with the lowest E-value was used. Sequences other than the V(D)J region were removed and transcripts containing incomplete V(D)J or/and frame shift were excluded. We then aligned each of the remaining transcripts to the assigned germline V gene using CLUSTALO^29^ and calculated the somatic hypermutation level.

To select representative antibodies for functional characterization, we first clustered all antibodies using USEARCH^30^ with the following criteria: identical heavy chain V and J gene assignments, the same length of CDRH3, and CDRH3 identity higher than 0.9. For each cluster, cells with the same light chain V and J gene assignments were grouped into a clone. All clone assignments were manually checked. We then calculated the clonal size for each clone, and one H and L chain pair per clone was chosen for antibody synthesis. For clones with multiple members, the member with the highest somatic hypermutation level was chosen for synthesis. For cells having multiple high quality H or L chains, which may be from doublets, we synthesized all H and L chain combinations.

### Analysis of S Trimer-Specific Antibody Repertoire

Because 88% of the S trimer-specific antibodies were IgG isotype, we therefore compared the repertoire features to IgG repertoires from three healthy donors^31^ (17,243 H chains, 27,575 kappa L chains, 20,889 lambda L chains). The repertoire data from the three healthy donors were combined and annotated using SONAR with the same process as above.

### Antibody Expression and Purification

For each antibody, variable genes were optimized for human cell expression and synthesized by GenScript. VH and VL were inserted separately into plasmids (gWiz or pcDNA3.4) that encode the constant region for H chain and L chain. Monoclonal antibodies were expressed in Expi293 (ThermoFisher, A14527) by co-transfection of H chain and L chain expressing plasmids using polyethylenimine and culture in 37°C shaker at 125 RPM and 8% CO_2_. On day 3 post transfection, 400 µL of supernatant were collected for screening for binding to S trimer and RBD by ELISA, and for neutralization of SARS-CoV-2 pseudovirus and authentic virus. Supernatants were also collected on day 5 for antibody purification by rProtein A Sepharose (GE, 17-1279-01) affinity chromatography.

### Production of Pseudoviruses

Recombinant Indiana VSV (rVSV) expressing SARS-CoV-2 spike was generated as previously described^32,33^. HEK293T cells were grown to 80% confluency before transfection with pCMV3-SARS-CoV-2-spike (Sino Biological) using FuGENE 6 (Promega). Cells were cultured overnight at 37°C with 5% CO_2_. The next day, medium was removed and VSV-G pseudotyped ΔG-luciferase (G*ΔG-luciferase, Kerafast) was used to infect the cells in DMEM at a MOI of 3 for 1 hr before washing the cells with 1X DPBS three times. DMEM supplemented with 2% fetal bovine serum and 100 I.U./mL of penicillin and 100 µg/mL of streptomycin were added to the inoculated cells, which were cultured overnight as described above. The supernatant was harvested the following day and clarified by centrifugation at 300 g for 10 mins before aliquoting and storing at −80°C.

### Pseudovirus Neutralization

Neutralization assays were performed by incubating pseudoviruses with serial dilutions of heat-inactivated plasma together with supernatant or purified antibodies, and scored by the reduction in luciferase gene expression. In brief, Vero E6 cells (ATCC) were seeded in a 96-well plate at a concentration of 2 × 10^4^ cells per well. Pseudoviruses were incubated the next day with serial dilutions of the test samples in duplicate or triplicate for 30 mins at 37°C. The mixture was added to cultured cells and incubated for an additional 24 hrs. The luminescence was measured by Britelite plus Reporter Gene Assay System (PerkinElmer). IC_50_ was defined as the dilution at which the relative light units were reduced by 50% compared with the virus control wells (virus + cells) after subtraction of the background in the control groups with cells only. The IC_50_ values were calculated using non-linear regression in GraphPad Prism 8.0.

### Authentic SARS-CoV-2 Neutralization

Supernatants containing expressed mAbs were diluted 1:10 and 1:50 in EMEM with 7.5% inactivated fetal calf serum and incubated with authentic SARS-CoV-2 (strain USA-WA1/2020; MOI 0.1) for 1hr at 37°C. Post-incubation, the mixture was transferred onto a monolayer of Vero-E6 cells that was cultured overnight. After incubation of the cells with the mixture for 70 hrs at 37°C, cytopathic effects (CPE) caused by the infection were scored for each well from 0 to 4 to indicate the degree of virus inhibition. Semi-quantitative representation of the inhibition for each antibody-containing supernatant at a dilution of 1:50 is shown in the lowest panel of Fig. 1b with neutralization levels ranging from (-) for none to (+++) for complete neutralization.

An end-point dilution assay in a 96-well plate format was performed to measure the neutralization activity of select purified mAbs. In brief, each antibody was serially diluted (5-fold dilutions) starting at 20 µg/mL. Triplicates of each mAb dilution were incubated with SARS-CoV-2 at a MOI of 0.1 in EMEM with 7.5% inactivated fetal calf serum for 1 hr at 37°C. Post incubation, the virus-antibody mixture was transferred onto a monolayer of Vero-E6 cells grown overnight. The cells were incubated with the mixture for 70 hrs. CPE were visually scored for each well in a blinded fashion by two independent observers. The results were then converted into percentage neutralization at a given mAb concentration, and the averages ± SEM were plotted using a five-parameter dose-response curve in GraphPad Prism 8.0.

### Epitope Mapping by ELISA

50 ng/well of S trimer, 50 ng/well of RBD, and 100 ng/well of NTD were coated on ELISA plates at 4°C overnight. The ELISA plates were then blocked with 300 μL of blocking buffer (1% BSA and 10% bovine calf serum (BCS) (Sigma) in PBS at 37°C for 2 hrs. Afterwards, supernatants from the antibody transfection or purified antibodies were serially diluted using dilution buffer (1% BSA and 20% BCS in PBS), incubated at 37°C for 1 hr. Next, 100 μL of 10,000-fold diluted Peroxidase AffiniPure goat anti-human IgG (H+L) antibody (Jackson ImmunoResearch) were added into each well and incubated for 1 hr at 37°C. The plates were washed between each step with PBST (0.5% Tween-20 in PBS). Finally, the TMB substrate (Sigma) was added and incubated before the reaction was stopped using 1M sulfuric acid. Absorbance was measured at 450 nm.

For the competition ELISA, purified mAbs were biotin-labeled using One-Step Antibody Biotinylation Kit (Miltenyi Biotec) following manufacturer recommendations and purified using 40K MWCO Desalting Column (ThermoFisher Scientific). 50 µL of serially diluted competitor antibodies were added into S trimer-precoated ELISA plates, followed by 50 µL of biotinylated antibodies at a concentration that achieves an OD_450_ reading of 1.5 in the absence of competitor antibodies. Plates were incubated at 37°C for 1 hr, and 100 µL of 500-fold diluted Avidin-HRP (ThermoFisher Scientific) were added into each well and incubated for another 1 hr at 37°C. The plates were washed by PBST between each of the previous steps. The plates were developed afterwards with TMB and absorbance was read at 450 nm after the reaction was stopped.

For the ACE2 competition ELISA, 100 ng of ACE2 protein (Abcam) was immobilized on the plates at 4°C overnight. The unbound ACE2 was washed away by PBST and then the plates were blocked. After washing, 100 ng of S trimer in 50 µL of dilution buffer was added into each well, followed by adding another 50 µL of serially diluted competitor antibodies and then incubating the plates at 37°C for 1 hr. The ELISA plates were washed 4 times by PBST and then 100 µL of 2000-fold diluted anti-strep-HRP (Millipore Sigma) were added into each well for another 1 hr at 37°C. The plates were then washed, developed with TMB, and absorbance was read at 450 nm after the reaction was stopped.

For all the competition ELISA experiments, the relative binding of biotinylated antibodies or ACE2 to the S trimer in the presence of competitors was normalized by comparing to competitor-free controls. Relative binding curve and the area under curve (AUC) were generated by fitting the non-linear five-parameter dose-response curve in GraphPad Prism 8.0.

### Cell-Surface Competition Binding Assay

Expi293 cells were co-transfected with vectors encoding pRRL-cPPT-PGK-GFP (Addgene) and pCMV3-SARS-CoV-2 (2019-nCoV) Spike (Sino Biological) at a ratio of 1:1. Two days after transfection, cells were incubated with a mixture of biotinylated mAb 2-43 (0.25 µg/mL) and serially diluted competitor antibodies at 4°C for 1 hr. Then 100 µL of diluted APC-streptavidin (Biolegend) were added to the cells and incubated at 4°C for 45 mins. Cells were washed 3 times with FACS buffer before each step. Finally, cells were resuspended and 2-43 binding to cell-surface S trimer was quantified on LSRII flow cytometer (BD Biosciences). The mean fluorescence intensity of APC in GFP-positive cells was analyzed using FlowJo and the relative binding of 2-43 to S trimer in the presence of competitors was calculated as the percentage of the mean fluorescence intensity compared to that of the competitor-free controls.

### Cryo-EM Data Collection and Processing

SARS-CoV-2 S trimer at a final concentration of 2 mg/ml was incubated with 6-fold molar excess per spike monomer of the antibody Fab fragments for 30 minutes in 10 mM sodium acetate pH 5.5, 150 mM NaCl, and 0.005% n-Dodecyl-β-D-maltoside (DDM). 2 µL of sample were incubated on C-flat 1.2/1.3 carbon grids for 30 secs and vitrified using a Leica EM GP Plunge Freezer. Data were collected on a Titan Krios electron microscope operating at 300 kV equipped with a Gatan K3 direct detector and energy filter using the Leginon software package^34^. A total electron fluence of 51.3 e/Å^2^ was fractionated over 40 frames, with a total exposure time of 2 secs. A magnification of 81,000x resulted in a pixel size of 1.058 Å, and a defocus range of -0.4 to -3.5 µm was used. All processing was done using cryoSPARC v2.14.2^35^. Raw movies were aligned and dose-weighted using patch motion correction, and the CTF was estimated using patch CTF estimation. A small subset of approximately 200 micrographs were picked using blob picker, followed by 2D classification and manual curation of particle picks, and used to train a Topaz neural network^36^. This network was then used to pick particles from the remaining micrographs, which were extracted with a box size of 384 pixels.

For the 2-4 Fab dataset, 2D classification followed by *ab initio* modelling and 3D heterogeneous refinement revealed 83,927 particles with three 2-4 Fabs bound, one to each RBD. A reconstruction of these particles using Non-Uniform Refinement with imposed C3 symmetry resulted in a 3.6 Å map, as determined by the gold standard FSC. Given the relatively low resolution of the RBD-Fab interface, masked local refinement was used to obtain a 3.5 Å map with significantly improved density. A masked local refinement of the remainder of the S timer resulted in a 3.5 Å reconstruction. These two local refinements were aligned and combined using the vop maximum function in UCSF Chimera^37^. This was repeated for the half maps, which were used, along with the refinement mask from the global Non-Uniform refinement, to calculate the 3DFSC^38^ and obtain an estimated resolution of 3.2 Å. All maps have been submitted to the EMDB with the ID EMD-22156.

For the 4-8 Fab dataset, image preprocessing and particle picking was performed as above. 2D classification, *ab initio* modelling, and 3D heterogeneous classification revealed 47,555 particles with 3 Fabs bound, one to each NTD and with all 3 RBDs in the down conformation. While this particle stack was refined to 3.9 Å using Non-Uniform refinement with imposed C3 symmetry, significant molecular motion prevented the visualization of the Fab epitope at high resolution (EMD-22159). In addition, 105,278 particles were shown to have 3 Fabs bound, but with 1 RBD in the up conformation. These particles were refined to 4.0 Å using Non-Uniform refinement with C1 symmetry (EMD-22158), and suffered from the same conformational flexibility as the all-RBD-down particles. This flexibility was visualized using 3D variability analysis in cryoSPARC.

For the 2-43 Fab dataset, which was collected at an electron fluence of 51.69 e/Å2, image preprocessing was performed as above, and particle picking was performed with blob picker. 2D classification, *ab initio* modelling, and 3D heterogeneous classification revealed 10,068 particles with 3 Fabs bound, which was refined to 5.8Å resolution (EMD-22157).

### Cryo-EM Model Fitting

An initial homology model of the 2-4 Fab was built using Schrodinger Release 2020-2: BioLuminate^39^. The RBD was initially modeled using the coordinates from PDB ID 6W41. The remainder of the S timer was modeled using the coordinates from PDB ID 6VSB. These models were docked into the consensus map using Chimera. The model was then fitted interactively using ISOLDE 1.0b5^40^ and COOT 0.8.9.2^41^, and using real space refinement in Phenix 1.18^42^. In cases where side chains were not visible in the experimental data, they were truncated to alanine. Validation was performed using Molprobity^43^ and EMRinger^44^. The model was submitted to the PDB with the ID 6XEY. Figures were prepared using ChimeraX^45^.

### Hamster Protection Experiment

*In vivo* evaluation of mAb 2-15 in an established golden Syrian hamster model of SARS-CoV-2 infection was performed as described previously with slight modifications^46^. Approval was obtained from the University of Hong Kong (HKU) Committee on the Use of Live Animals in Teaching and Research. Briefly, 6-8-week-old male and female hamsters were obtained from the Chinese University of Hong Kong Laboratory Animal Service Centre through the HKU Laboratory Animal Unit and kept in Biosafety Level-2 (BSL-2) housing with access to standard pellet feed and water *ad libitum* until virus challenge in the BSL-3 animal facility. Each hamster (n = 4 per group) was intraperitoneally administered one dose of 1.5 mg/kg of mAb 2-15 in phosphate-buffered saline (PBS), 0.3 mg/kg of mAb 2-15 in PBS, or PBS alone as control. Twenty-four hours later, each hamster was intranasally inoculated with a challenge dose of 100 µL of Dulbecco’s Modified Eagle Medium containing 10^5^ PFU of SARS-CoV-2 (HKU-001a strain, GenBank accession no: MT230904.1) under intraperitoneal ketamine (200 mg/kg) and xylazine (10 mg/kg) anesthesia. The hamsters were monitored twice daily for clinical signs of disease and sacrificed at day 4 post-challenge. Half of each hamster’s lung tissues were used for viral load determination by a quantitative SARS-CoV-2 RdRp/Hel reverse transcription-polymerase chain reaction assay^47^ and an infectious virus titration using a plaque assay described previously^46^. Student’s *t* test was used to determine significant differences among the groups, and a p value of < 0.05 was considered statistically significant.

### Data Availability

The 19 neutralizing antibodies were deposited to Genbank with accession numbers from MT712278 to MT712315. Coordinates for the antibody 2-4 complex are deposited in the Protein Data Bank as PDB 6XEY. Cryo-EM maps and data are deposited in EMDB with deposition codes EMDB-22156 for antibody 2-4, EMDB-22158 and EMDB-22159 for antibody 4-8, and EMDB-22275 for antibody 2-43. These data are used in Fig. 4 and Extended Data Figs. 7, 8, 9, 10, and 11.

### Code Availability

Next-generation sequencing data of antibody repertoires were processed using Cell ranger v3.1.0, SONAR V1, BLAST v2.2.25, CLUSTALO1.2.3, and USEARCH v9.2.64. Cryo-EM data was collected using Leginon 3.4.beta. Cryo-EM data was processed using cryoSPARC v2.14.2, MotionCor2, Topaz v0.2.4, 3DFSC v3.0, UCSF Chimera v1.13.1, ChimeraX v0.93, ISOLDE v1.0b5, Phenix v1.18, and COOT v0.8.9.2.

## Acknowledgements

We thank N. Wang and J. McLellan for providing reagents for generating the SARS-CoV-2 S trimer and RBD-SD1 with mFc tag. We thank W. Chen for assistance with generating the Venn diagrams, and B. DeKosky and X. Wu for helpful input. This study was supported by funding to D.D.H. from the Jack Ma Foundation, the JPB Foundation, Samuel Yin, Brii Biosciences, Tencent Charity Foundation, Roger Wu, Carol Ludwig, and Peggy and Andrew Cherng. Cryo-EM data collection was performed at the National Center for CryoEM Access and Training and the Simons Electron Microscopy Center located at the New York Structural Biology Center, supported by the NIH Common Fund Transformative High Resolution Cryo-Electron Microscopy program (U24 GM129539) and by grants from the Simons Foundation (SF349247) and NY State Assembly. Data analysis was performed at the National Resource for Automated Molecular Microscopy (NRAMM), supported by the NIH National Institute of General Medical Sciences (GM103310). Hamster experiments were conducted with support from the Health@InnoHK (Centre for Virology, Vaccinology and Therapeutics), Innovation and Technology Commission, The Government of the Hong Kong Special Administrative Region.

## Author Contributions

D.D.H conceived of the project. L.L., P.W., M.S.N, J.Y., Q.W., Y.H. performed many of the experiments. M.T.Y. was responsible for recruiting patients, obtaining clinical specimens, and summarizing clinical data. L.L., V.S., A.F. and X.V.G. performed and analyzed the B-cell sorting, 10X Genomics, sequencing and analysis of the clones. Z.S. performed bioinformatic analyses on 10X next-generation sequencing data and antibody repertoire. J.Y. cloned, expressed, and purified the mAbs. L.L. and Q.W. performed the epitope mapping and binding experiments. P.W. conducted the pseudovirus neutralization assays and M.S.N. and Y.X. performed infectious SARS-CoV-2 neutralization assays. M.A.R., G.C., J.B, J.G., and L.S. carried out the cryo-EM studies. J.F-W.C., Z.C., and K-Y.Y. were responsible for the hamster experiment. Y.L. helped with project management. T.Z. and P.D.K. provided key reagents for the study, and P.D.K. contributed to the analysis and discussion of the data. L.L., P.W., M.S.N., J.Y., Y.H., Z.S., M.A.R., Q.W, L.S., and D.D.H analyzed the results, and D.D.H. wrote the manuscript, with contributions from each author. J.G.S. provided valuable suggestions.

## Ethic Statement

Acquiring convalescent specimens to isolate and identify potent monoclonal antibodies against COVID-19” (AAAS9517) was approved by the Columbia University Institutional Review Board. Informed consent was obtained from all participants or surrogates.

## Competing Interest Declaration

A provisional patent application has been filed for the monoclonal antibodies described in the manuscript. L.L. and D.D.H. are inventors.

## Extended Data

**Extended Data Table 1.** Patient information.

| Patient | Age | Sex & Race | Days from symptom onset to: | Biomarker | Complications | Outcome |
| --- | --- | --- | --- | --- | --- | --- |
| 1 | 57 | Female, Hispanic | Admission: 7<br>MV: 12<br>Ab isolation: 18 | hsCRP = 208 mg/L<br>ESR = 58 mm/hr<br>IL-6 = 23 pg/mL<br>Ferritin = 766 ng/mL<br>D-dimer = 3.4 µg/mL<br>FEU | ARDS | Discharged on day 30 |
| 2 | 71 | Female, Hispanic | Admission: 20<br>MV: 20<br>Ab isolation: 29 | hsCRP = 33 mg/L<br>ESR > 130 mm/hr<br>IL-6 = 13 pg/mL<br>Ferritin = 425 ng/mL<br>D-dimer = 5.7 µg/mL<br>FEU | ARDS<br>Ventilator associated pneumonia | Discharged on day 45 |
| 3 | 61 | Male, White | Admission: 10<br>MV: 10<br>Ab isolation: 21 | hsCRP = 51 mg/L<br>ESR = 57 mm/hr<br>IL-6 > 315 pg/mL<br>Ferritin = 3,238 ng/mL<br>D-dimer = 7.4 µg/mL<br>FEU | ARDS<br>Acute kidney injury (hemodialysis)<br>Sepsis | Death on day 28 |
| 4 | 51 | Male, Black | Admission: 7<br>MV: 10<br>Ab isolation: 25 | hsCRP = 88 mg/L<br>ESR = 110 mm/hr<br>IL-6 = 77 pg/mL<br>Ferritin = 510 ng/mL<br>D-dimer = 13.4 µg/mL<br>FEU | ARDS<br>Acute kidney injury (no hemodialysis)<br>Ventilator associated pneumonia | Discharged on day 51 |
| 5 | 50 | Male, White | Admission: 5<br>MV: 7<br>Ab isolation: 32 | hsCRP = 2 mg/L<br>ESR = 63 mm/hr | ARDS<br>Neuropathy | Discharged on day 27 |

Abbreviation: ARDS, acute respiratory distress syndrome; MV, mechanical ventilation; hsCRP,
high sensitivity C-reactive protein, ULN>10 mg/L; ESR, erythrocyte sedimentation rate,
ULN=20 mm/hr; Interleukin 6, ULN= 5 pg/mL; Ferritin, ULN=150 ng/mL; D-dimer quantitative
ULN= 0.8ug/mL FEU.

**Extended Data Table 2.** Cryo-EM data collection, refinement, and validation statistics.

|  | SARS-CoV-2 spike with<br>Fab 2-4<br>(EMDB-22156)<br>(PDB 6XEY) | SARS-CoV-2 spike<br>RBD up with Fab 4-8<br>(EMDB-22158) | SARS-CoV-2 spike<br>RBD down with<br>Fab 4-8<br>(EMDB-22159) | SARS-CoV-2 spike<br>with Fab 2-43<br>(EMDB-22275) |
| --- | --- | --- | --- | --- |
| **Data collection and processing** |  |  |  |  |
| Magnification | 81,000 | 81,000 | 81,000 | 81,000 |
| Voltage (kV) | 300 | 300 | 300 | 300 |
| Electron exposure (e-/Å<sup>2</sup>) | 51.30 | 51.30 | 51.30 | 51.69 |
| Defocus range (μm) | -0.4 to -3.5 | -0.4 to -3.5 | -0.4 to -3.5 | -0.4 to -3.5 |
| Pixel size (Å) | 1.058 | 1.058 | 1.058 | 1.058 |
| Symmetry imposed | C3 | C1 | C3 | C1 |
| Initial particle images (no.) | 556,983 | 256,848 | 256,848 | 55,161 |
| Final particle images (no.) | 83,927 | 105,278 | 47,555 | 10,068 |
| Map resolution (Å) | 3.25 | 4.0 | 3.9 | 5.8 |
| FSC threshold | 0.143 | 0.143 | 0.143 | 0.143 |
| Map resolution range (Å) | 406.3-3.25 | 406.3-4.0 | 406.3-3.9 | 406.3-5.8 |
| **Refinement** |  |  |  |  |
| Initial model used (PDB code) | 6VSB |  |  |  |
| Model resolution (Å) | 3.7 |  |  |  |
| FSC threshold | 0.5 |  |  |  |
| Model resolution range (Å) | 406.3-3.25 |  |  |  |
| Map sharpening *B* factor (Å<sup>2</sup>) | -97.5 |  |  |  |
| Model composition |  |  |  |  |
| Non-hydrogen atoms | 28,482 |  |  |  |
| Protein residues | 3788 |  |  |  |
| Ligands | 63 |  |  |  |
| *B* factors (Å<sup>2</sup>) |  |  |  |  |
| Protein | 54.35 |  |  |  |
| Ligand | 73.91 |  |  |  |
| R.m.s. deviations |  |  |  |  |
| Bond lengths (Å) | 0.005 |  |  |  |
| Bond angles (°) | 0.810 |  |  |  |
| Validation |  |  |  |  |
| MolProbity score | 1.51 |  |  |  |
| Clashscore | 3.59 |  |  |  |
| Poor rotamers (%) | 0.22 |  |  |  |
| Ramachandran plot |  |  |  |  |
| Favored (%) | 94.75 |  |  |  |
| Allowed (%) | 5.17 |  |  |  |
| Disallowed (%) | 0.08 |  |  |  |

**Extended Data Fig. 1.**
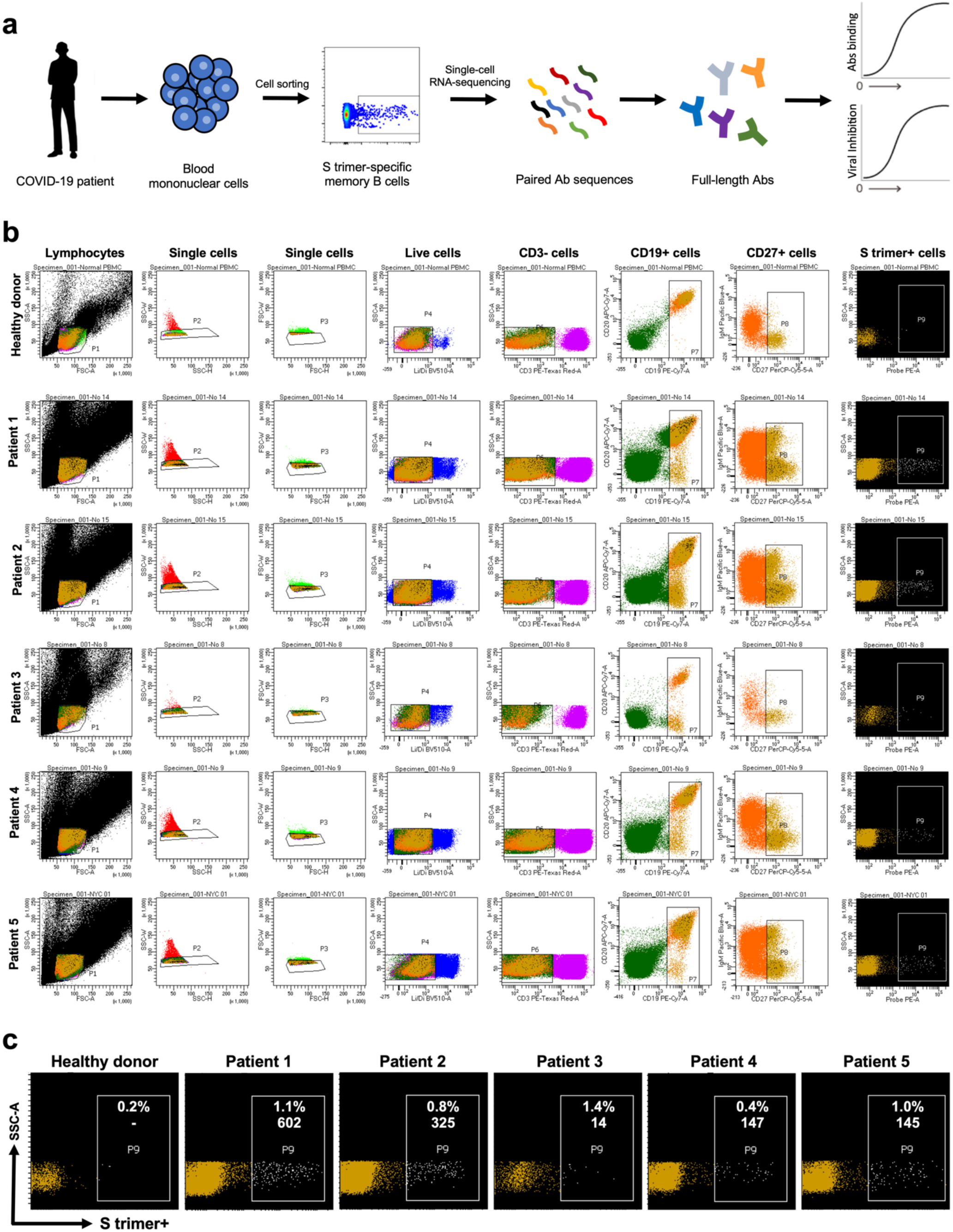
SARS-CoV-2 S trimer-specific antibody isolation strategy. **a**, Schema for isolating of S trimer-specific mAbs from memory B cells in the blood of infected patients. **b**, Sorting results on the isolation of S trimer-specific memory B cells using flow cytometry. **c**, Magnified representation of the panel of S trimer-positive memory B cells for each patient. Inset numbers indicate the absolute number and the percentage of S trimer-specific memory B cells isolated from each case.

**Extended Data Fig. 2.**
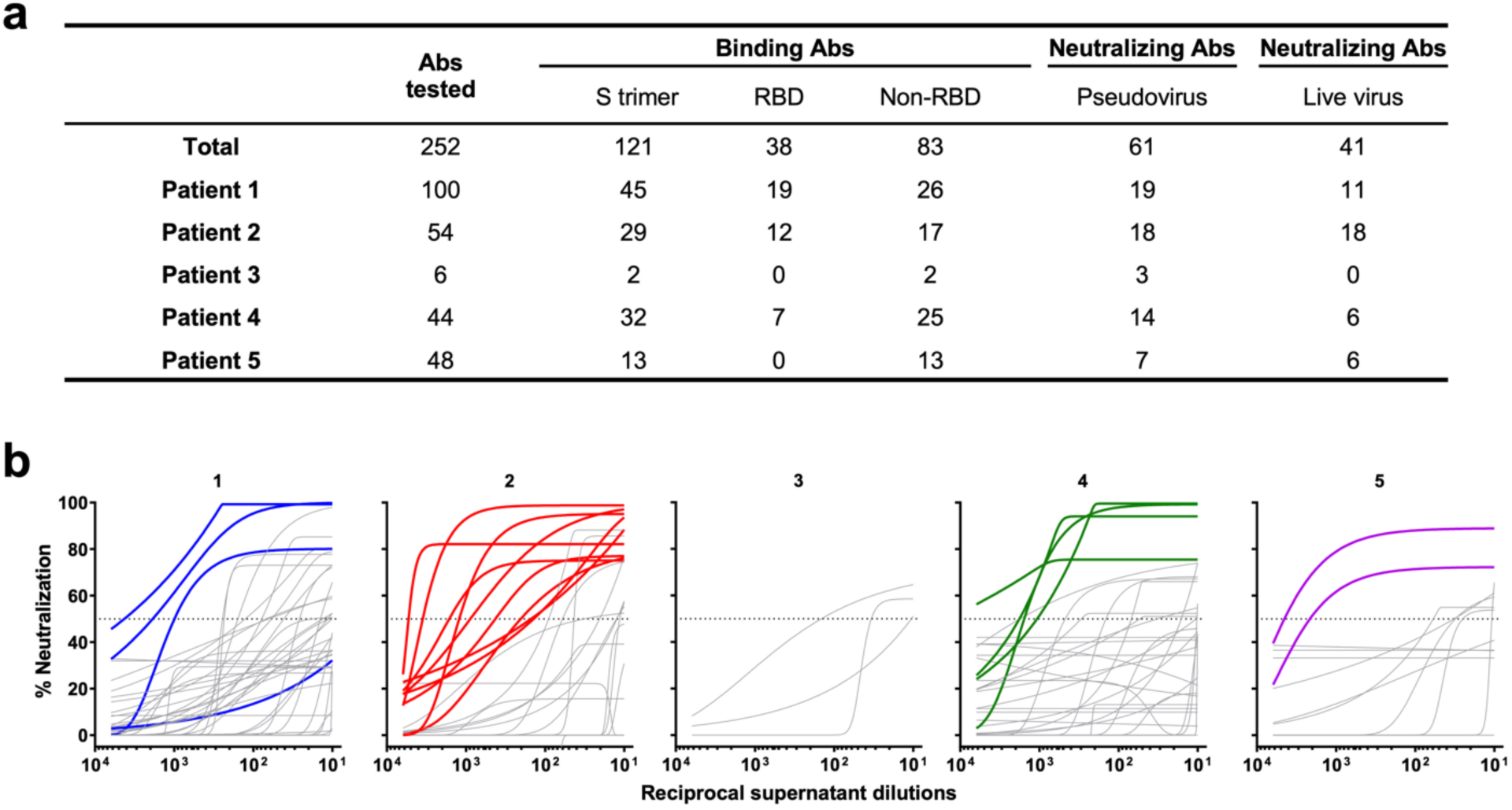
Summary of mAb screening of transfection supernatants. **a**, Numbers of binding and neutralizing antibodies from patients 1 to 5. **b**, The best-fit pseudovirus neutralization curves for 130 samples that were positive in at least one of the screens shown in Fig. 1b. The 18 transfection supernatants that showed evidently better potency are highlighted in colors, while others with non-neutralizing or weakly neutralizing activities are shown in grey. One additional supernatant (Patient 1) that was initially missed in the pseudovirus screen but later found to be a potent neutralizing mAb (1-87) is also highlighted.

**Extended Data Fig. 3.**
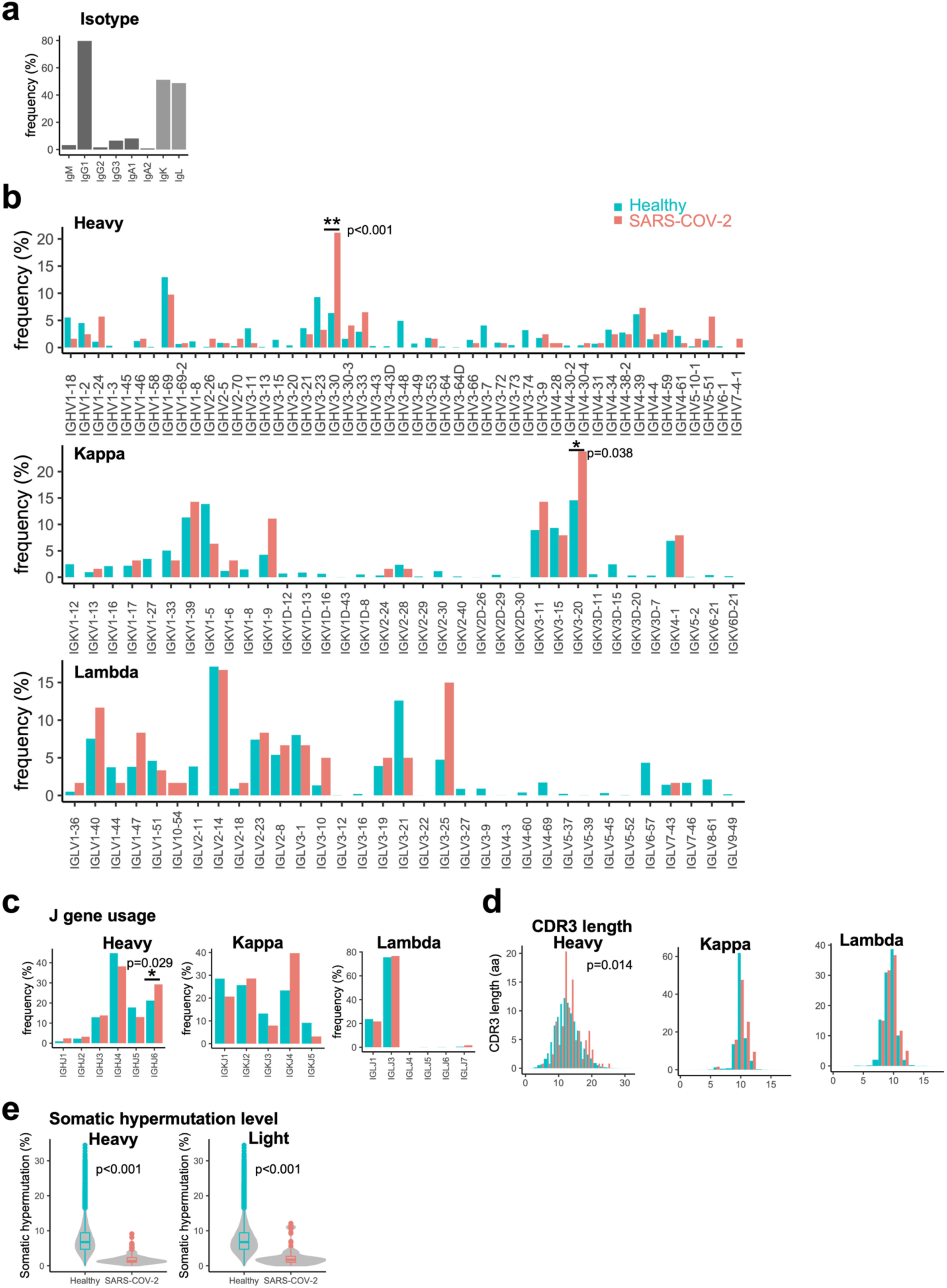
Genetic features of SARS-CoV-2-specific antibody repertoire. **a**, 108 of the 123 antigen-specific antibodies are from IgG isotype. The kappa and lambda light chains are comparably used. **b**, Compared to IgG repertoires of healthy human donors (17,243, 27,575, and 20,889 transcripts for heavy, kappa, and lambda chains respectively), IGHV3-30 (antigen-specific n=26 and healthy donor n=1117) and IGKV3-20 genes (antigen-specific n=15 and healthy donor n=4,071) are over-represented in heavy and light chain repertoires respectively (P values are 6.415e-11 and 0.04332 respectively, χ2-test with 1 degree of freedom). We did not test the enrichment of other genes because the numbers of antigen-specific antibodies are less than 15. **c**, The usage of IGHJ6 gene (antigen-specific n=36 and healthy donor n=3646) was significantly higher in antigen-specific antibodies (χ2-test with 1 degree of freedom, p=0.02807). **d**, The CDRH3 length of antigen-specific antibodies is significantly longer than in healthy donors (two-sided Kolmogorov–Smirnov test, p=0.014). **e**, For both heavy and light chains, the V region nucleotide somatic hypermutation levels are significantly lower than in antibodies of healthy donors (two-sided Kolmogorov–Smirnov test, p< 2.2e-16 for both heavy and light chains). For the boxplots, the middle lines are medians. The lower and upper hinges correspond to the first and third quartiles respectively. The upper whisker extends to values no larger than 1.5*IQR (the inter-quartile range or distance between the first and third quartiles) from the hinge. The lower whisker extends to values no smaller than 1.5 * IQR from the hinge. Data points beyond the whiskers were plotted as outliers using dots.

**Extended Data Fig. 4.**
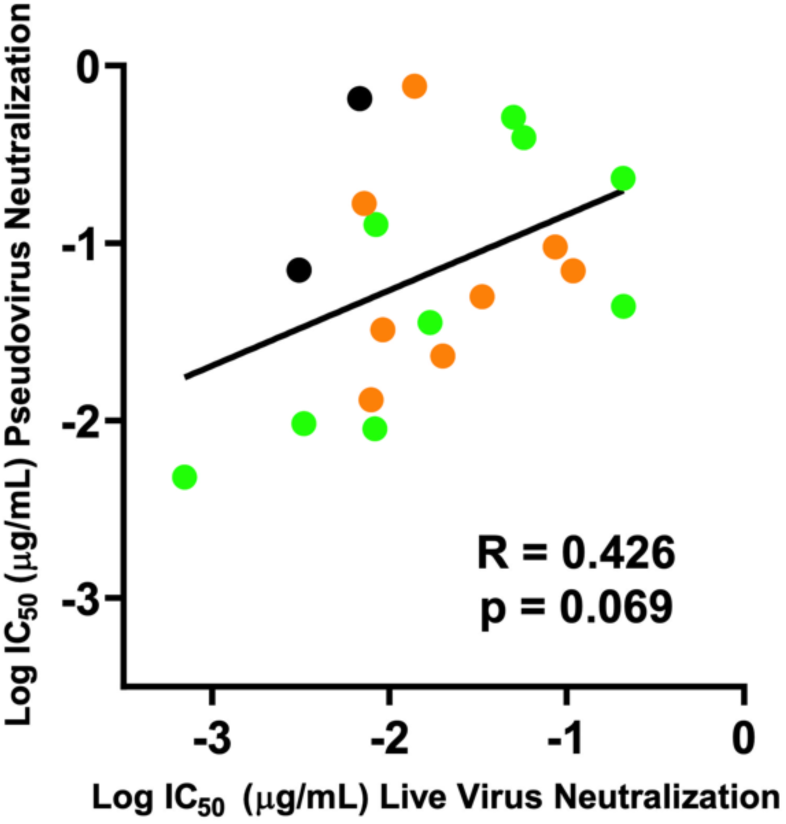
Correlation of neutralizing antibody titers of the top 19 mAbs in the live SARS-CoV-2 assay versus the pseudovirus assay. Green circles represent RBD-directed antibodies; orange circles represent NTD-directed antibodies; and black circles represent antibodies in the “Others” category. The *Pearson correlation coefficient (r)* and the *p* value were calculated using GraphPad Prism. Experiments were performed in triplicates for all mAbs tested.

**Extended Data Fig. 5.**
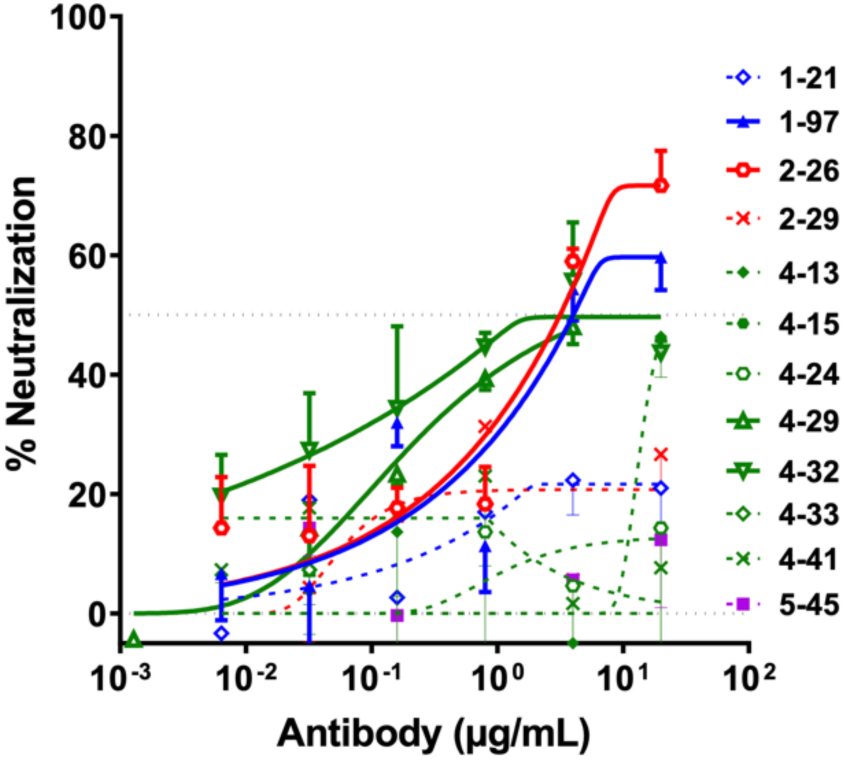
The pseudovirus neutralization profiles for 12 purified mAbs that strongly bound the S trimer but with weak or no virus-neutralizing activities. The four mAbs with weak neutralizing activities against SARS-CoV-2 pseudovirus are shown in sold lines, and the remaining 8 non-neutralizing mAbs are shown in dashed lines.

**Extended Data Fig. 6.**
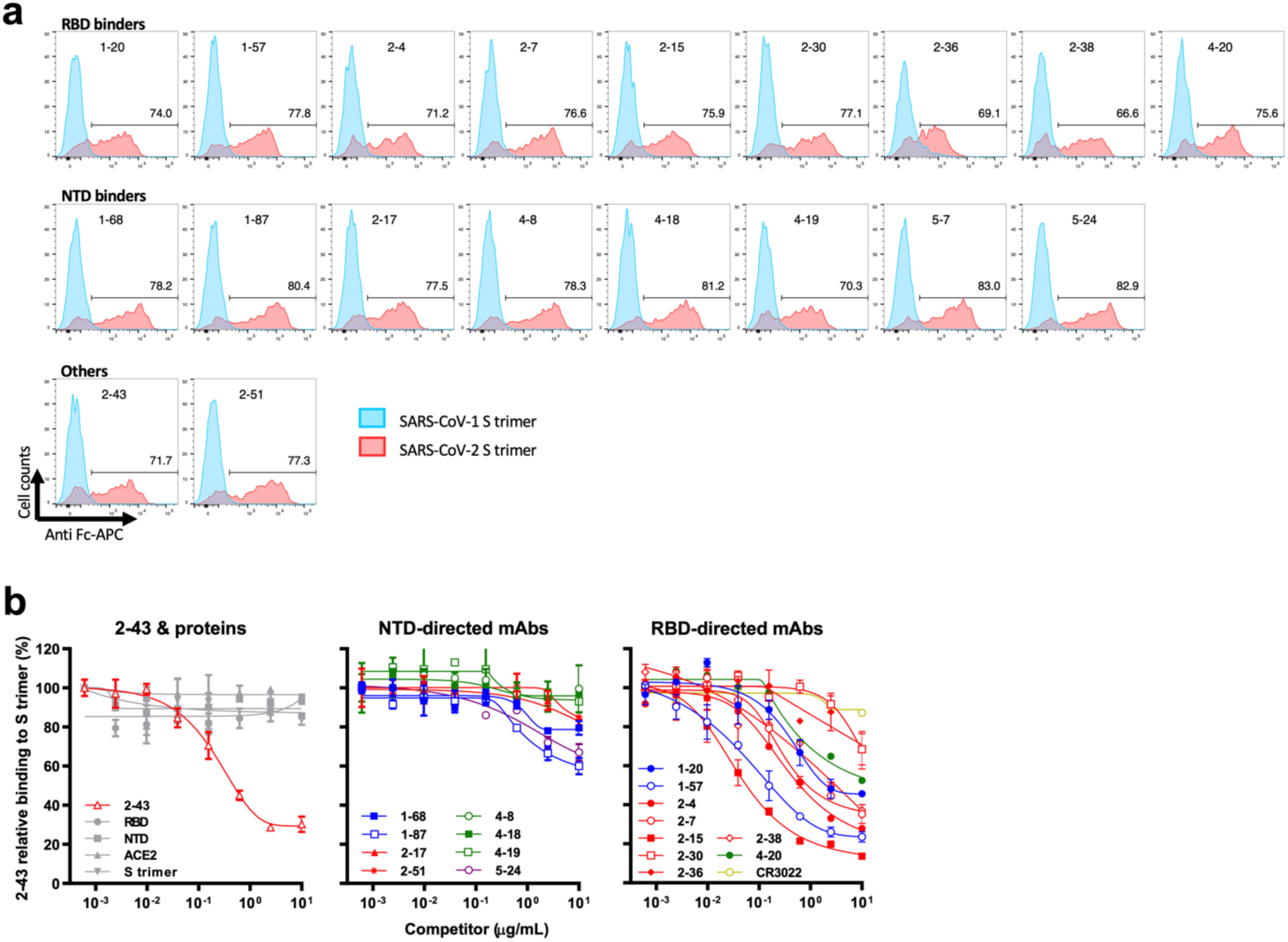
Cell-surface staining with antibodies. **a**, Antibody binding to the SARS-CoV-1 (blue) and SARS-CoV-2 (red) spike proteins expressed on the cell surface. Expi293 cells were co-transfected with GFP and full-length SARS-CoV-1 or SARS-CoV-2 spike genes. After 48 hours, antibody binding to spike protein in the GFP-positive cells was detected by flow cytometer. The data show all antibodies tested were able to recognize the wildtype SARS-CoV-2 spike protein but not SARS-CoV-1 spike protein. **b**, Monoclonal Ab 2-43 bound to S trimer expressed on Expi293 cell surface can be competed out by mAbs directed to RBD but only minimally by mAbs to the NTD region. Shown are representative data from three independent experiments.

**Extended Data Fig. 7.**
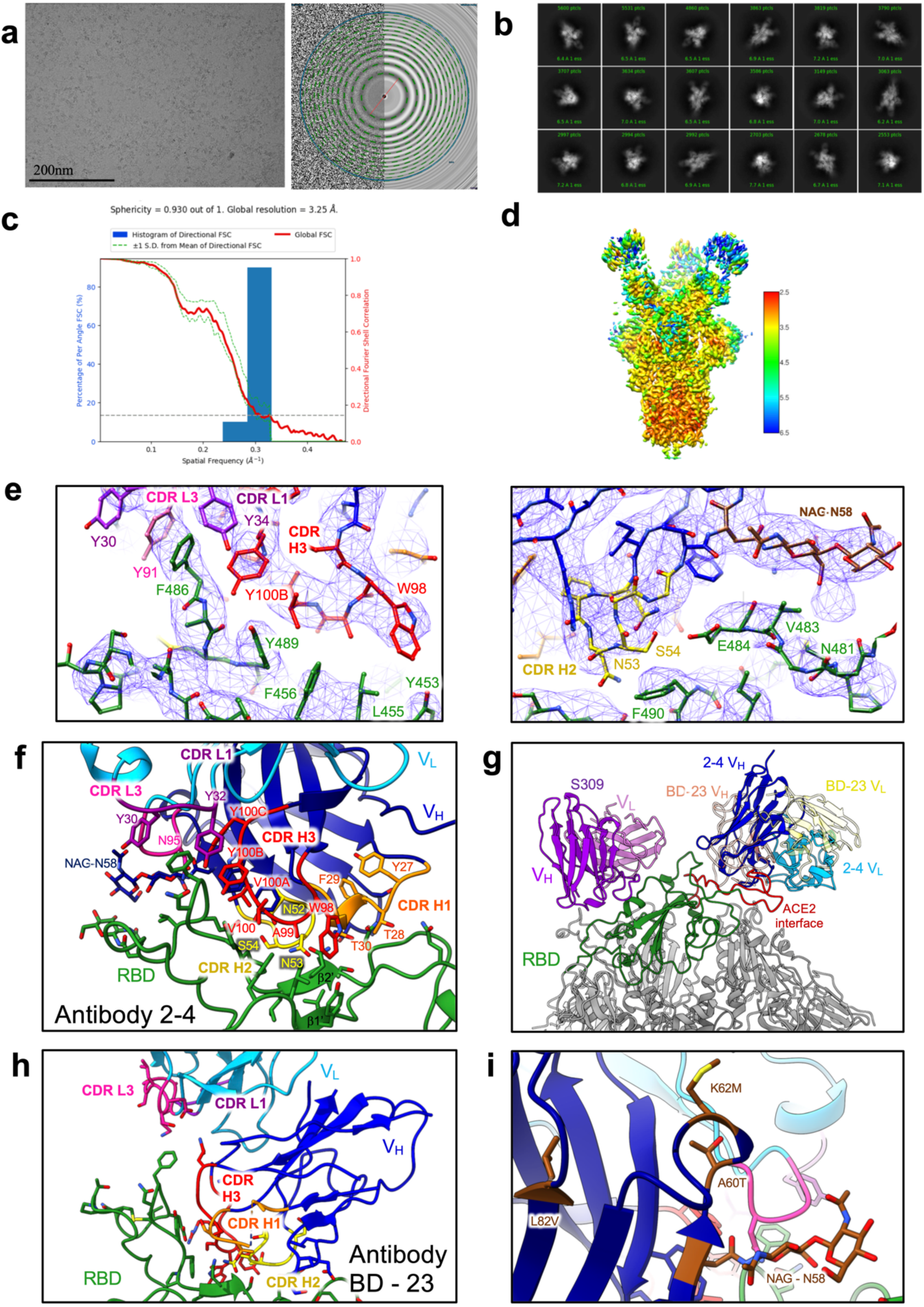
Cryo-EM analysis of antibody 2-4 in complex with the S trimer. **a**, Representative micrograph and CTF of the micrograph. 8,324 micrographs were collected in total. **b**, Representative 2D class averages. **c**, Resolution of the consensus map with C3 symmetry as calculated by 3DFSC. **d**, The local resolution of the full map as calculated by cryoSPARC at an FSC cutoff of 0.5. **e**, Representative density of the Fab 2-4 (blue) and RBD (green) interface, showing interactions of CDR H3 in red, L1 in magenta, and L3 in light magenta (left), along with CDR H2 and the N-linked glycosylation added by SHM at ASN58 (right). **f**, Fab 2-4 binding interface with RBD. V_H_ is shown in blue, V_L_ in light blue, with CDRs H1 in orange, H2 in yellow, H3 in red, L1 in magenta, and L3 in light magenta. **g**, Positions of antibodies 2-4, S309^8^, and BD-23^9^ on the trimeric CoV-2 spike. **h**, Antibody BD-23^9^ in complex with S trimer. **i**, Somatic hypermutations found only in the antibody 2-4 heavy chain, shown in brown. The mutation A60T creates an NxT sequence leading to N58 glycosylation.

**Extended Data Fig. 8.**
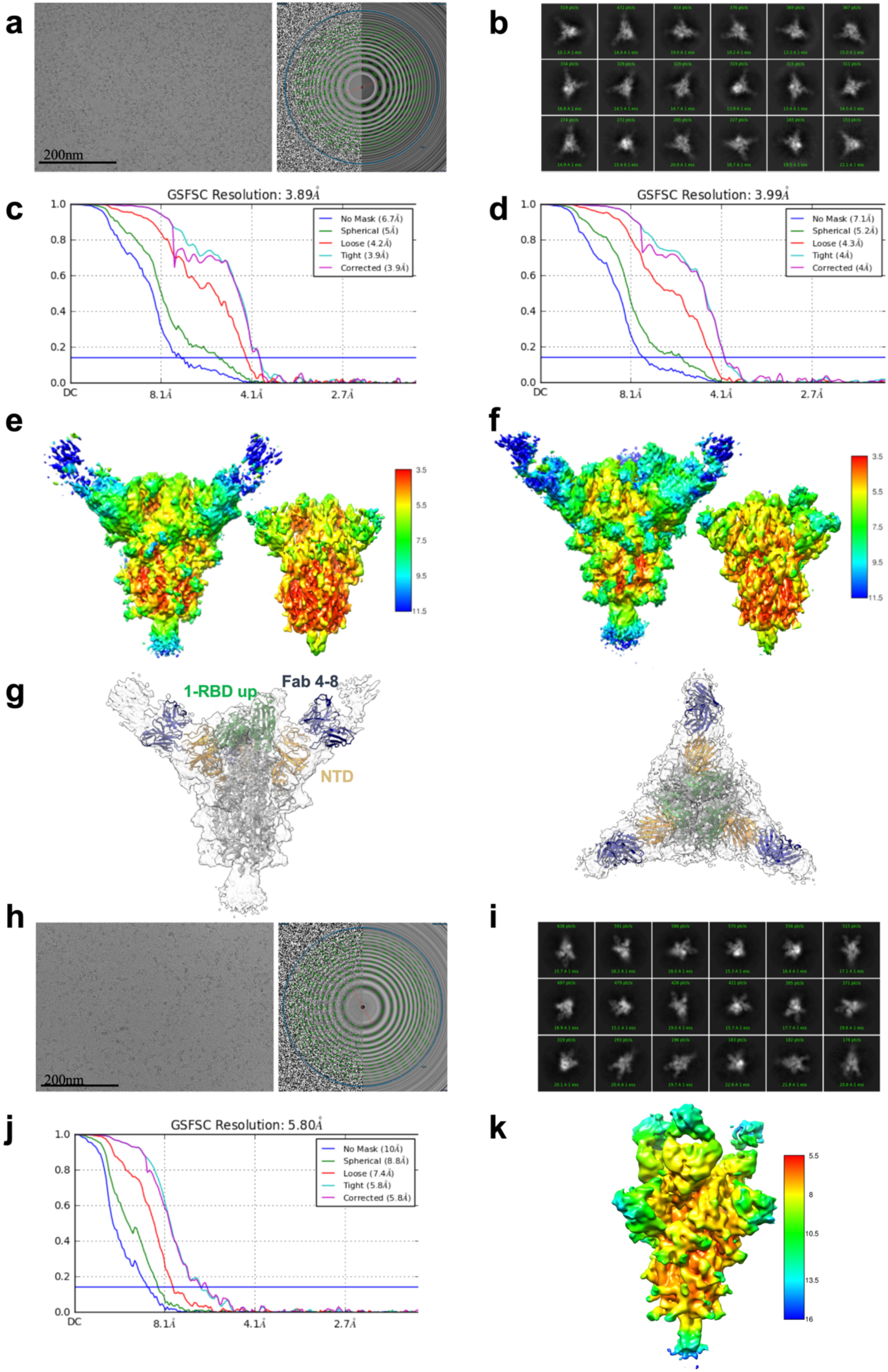
Cryo-EM data processing for antibodies 4-8 and 2-43 in complex with S trimer. **a**, Representative 4-8 micrograph and CTF of the micrograph. 3,153 micrographs were collected in total. **b**, Representative 2D class averages. **c**, Resolution of the spike in the RBD down conformation in complex with Fab 4-8. **d**, Resolution of the spike in the RBD up conformation in complex with Fab 4-8. **e**, Local resolution of the spike in the RBD down conformation in complex with Fab 4-8 at an FSC cutoff of 0.5, with two thresholds shown. **f**, Local resolution of the spike in the RBD up conformation in complex with Fab 4-8 at an FSC cutoff of 0.5, with two thresholds shown. **g**, Although the map was reconstructed at 4.0Å resolution, density for 4-8 Fab is poor due to molecular motion. A rigid body fit with SARS-CoV-2 spike and an antibody variable domain model is shown. **h-k**, Cryo-EM data processing for antibody 2-43 in complex with the S trimer. **h**, Representative 2-43 micrograph and CTF of the micrograph. **i**, Representative 2D class averages. **j**, Resolution of Fab 2-43 in complex with S trimer. **k**, The local resolution of the full map as calculated by cryoSPARC at an FSC cutoff of 0.5.

## References

1 Zhou, P. et al. A pneumonia outbreak associated with a new coronavirus of probable bat origin. Nature 579, 270–273, doi: 10.1038/s41586-020-2012-7 (2020).

2 Wang, C., Horby, P. W., Hayden, F. G. & Gao, G. F. A novel coronavirus outbreak of global health concern. Lancet 395, 470–473, doi: 10.1016/S0140-6736(20)30185-9 (2020).

3 Walls, A. C. et al. Structure, Function, and Antigenicity of the SARS-CoV-2 Spike Glycoprotein. Cell 181, 281–292 e286, doi: 10.1016/j.cell.2020.02.058 (2020).

4 Wrapp, D. et al. Cryo-EM structure of the 2019-nCoV spike in the prefusion conformation. Science 367, 1260–1263, doi: 10.1126/science.abb2507 (2020).

5 Hoffmann, M. et al. SARS-CoV-2 Cell Entry Depends on ACE2 and TMPRSS2 and Is Blocked by a Clinically Proven Protease Inhibitor. Cell 181, 271–280 e278, doi: 10.1016/j.cell.2020.02.052 (2020).

6 Wang, Q. et al. Structural and Functional Basis of SARS-CoV-2 Entry by Using Human ACE2. Cell 181, 894–904 e899, doi: 10.1016/j.cell.2020.03.045 (2020).

7 Ju, B. et al. Human neutralizing antibodies elicited by SARS-CoV-2 infection. Nature, doi: 10.1038/s41586-020-2380-z (2020).

8 Pinto, D. et al. Cross-neutralization of SARS-CoV-2 by a human monoclonal SARS-CoV antibody. Nature, doi: 10.1038/s41586-020-2349-y (2020).

9 Cao, Y. et al. Potent neutralizing antibodies against SARS-CoV-2 identified by high-throughput single-cell sequencing of convalescent patients’ B cells. Cell, doi: 10.1016/j.cell.2020.05.025 (2020).

10 Wu, Y. et al. A noncompeting pair of human neutralizing antibodies block COVID-19 virus binding to its receptor ACE2. Science 368, 1274–1278, doi: 10.1126/science.abc2241 (2020).

11 Hansen, J. et al. High-Throughput Effort Using Both Humanized Mice and Convalescent Humans Yields SARS-CoV-2 Antibody Cocktail. Science, doi: in press (2020).

12 Sheng, Z. et al. Gene-Specific Substitution Profiles Describe the Types and Frequencies of Amino Acid Changes during Antibody Somatic Hypermutation. Front Immunol 8, 537, doi: 10.3389/fimmu.2017.00537 (2017).

13 ter Meulen, J. et al. Human monoclonal antibody combination against SARS coronavirus: synergy and coverage of escape mutants. Plos Med 3, e237, doi: 10.1371/journal.pmed.0030237 (2006).

14 Tian, X. et al. Potent binding of 2019 novel coronavirus spike protein by a SARS coronavirus-specific human monoclonal antibody. Emerg Microbes Infect 9, 382–385, doi: 10.1080/22221751.2020.1729069 (2020).

15 Yuan, M. et al. A highly conserved cryptic epitope in the receptor binding domains of SARS-CoV-2 and SARS-CoV. Science 368, 630–633, doi: 10.1126/science.abb7269 (2020).

16 Rogers, T. F. et al. Rapid isolation of potent SARS-CoV-2 neutralizing antibodies and protection in a small animal model. bioRxiv, doi: 10.1101/2020.05.11.088674 (2020).

17 Chen, X. et al. Human monoclonal antibodies block the binding of SARS-CoV-2 spike protein to angiotensin converting enzyme 2 receptor. Cell Mol Immunol 17, 647–649, doi: 10.1038/s41423-020-0426-7 (2020).

18 Zeng, X. et al. Isolation of a human monoclonal antibody specific for the receptor binding domain of SARS-CoV-2 using a competitive phage biopanning strategy. Antibody Therapeutics 3, 95–100, doi: 10.1093/abt/tbaa008 (2020).

19 Liu, X. et al. Neutralizing Antibodies Isolated by a site-directed Screening have Potent Protection on SARS-CoV-2 Infection. bioRxiv, 2020.2005.2003.074914, doi: 10.1101/2020.05.03.074914 (2020).

20 Zost, S. J. et al. Rapid isolation and profiling of a diverse panel of human monoclonal antibodies targeting the SARS-CoV-2 spike protein. bioRxiv, doi: 10.1101/2020.05.12.091462 (2020).

21 Robbiani, D. F. et al. Convergent Antibody Responses to SARS-CoV-2 Infectijon in Convalescent Individuals. bioRxiv, doi: 10.1101/2020.05.13.092619 (2020).

22 Brouwer, P. J. M. et al. Potent neutralizing antibodies from COVID-19 patients define multiple targets of vulnerability. bioRxiv, 2020.2005.2012.088716, doi: 10.1101/2020.05.12.088716 (2020).

23 Chi, X. et al. A potent neutralizing human antibody reveals the N-terminal domain of the Spike protein of SARS-CoV-2 as a site of vulnerability. bioRxiv, 2020.2005.2008.083964, doi: 10.1101/2020.05.08.083964 (2020).

24 Wang, C. et al. A human monoclonal antibody blocking SARS-CoV-2 infection. Nat Commun 11, 2251, doi: 10.1038/s41467-020-16256-y (2020).

25 Wang, P. et al. SARS-CoV-2 Neutralizing Antibody Responses Are More Robust in Patients with Severe Disease. bioRxiv, 2020.2006.2013.150250, doi: 10.1101/2020.06.13.150250 (2020).

## Methods references

26 Schramm, C. A. et al. SONAR: A High-Throughput Pipeline for Inferring Antibody Ontogenies from Longitudinal Sequencing of B Cell Transcripts. Frontiers in immunology 7, 372, doi:10.3389/fimmu.2016.00372 (2016).

27 Altschul, S. F. et al. Gapped BLAST and PSI-BLAST: a new generation of protein database search programs. Nucleic Acids Res 25, 3389–3402 (1997).

28 Lefranc, M. P. et al. IMGT, the international ImMunoGeneTics information system. Nucleic Acids Res 37, D1006–1012, doi:10.1093/nar/gkn838 (2009).

29 Sievers, F. & Higgins, D. G. Clustal Omega, Accurate Alignment of Very Large Numbers of Sequences. Multiple Sequence Alignment Methods 1079, 105–116, doi:10.1007/978-1-62703-646-7_6 (2014).

30 Edgar, R. C. Search and clustering orders of magnitude faster than BLAST. Bioinformatics 26, 2460–2461, doi:10.1093/bioinformatics/btq461 (2010).

31 Sheng, Z. et al. Gene-Specific Substitution Profiles Describe the Types and Frequencies of Amino Acid Changes during Antibody Somatic Hypermutation. Frontiers in immunology 8, 537 (2017).

32 Nie, J. et al. Establishment and validation of a pseudovirus neutralization assay for SARS-CoV-2. Emerg Microbes Infect 9, 680–686, doi:10.1080/22221751.2020.1743767 (2020).

33 Whitt, M. A. Generation of VSV pseudotypes using recombinant DeltaG-VSV for studies on virus entry, identification of entry inhibitors, and immune responses to vaccines. J Virol Methods 169, 365–374, doi:10.1016/j.jviromet.2010.08.006 (2010).

34 Suloway, C. et al. Automated molecular microscopy: the new Leginon system. J Struct Biol 151, 41–60, doi:10.1016/j.jsb.2005.03.010 (2005).

35 Punjani, A., Rubinstein, J. L., Fleet, D. J. & Brubaker, M. A. cryoSPARC: algorithms for rapid unsupervised cryo-EM structure determination. Nat Methods 14, 290–296, doi:10.1038/nmeth.4169 (2017).

36 Bepler, T. et al. Positive-unlabeled convolutional neural networks for particle picking in cryo-electron micrographs. Nature Methods 16, 1153–1160, doi:10.1038/s41592-019-0575-8 (2019).

37 Pettersen, E. F. et al. UCSF Chimera--a visualization system for exploratory research and analysis. J Comput Chem 25, 1605–1612, doi:10.1002/jcc.20084 (2004).

38 Tan, Y. Z. et al. Addressing preferred specimen orientation in single-particle cryo-EM through tilting. Nat Methods 14, 793–796, doi:10.1038/nmeth.4347 (2017).

39 Zhu, K. et al. Antibody structure determination using a combination of homology modeling, energy-based refinement, and loop prediction. Proteins 82, 1646–1655, doi:10.1002/prot.24551 (2014).

40 Croll, T. I. ISOLDE: a physically realistic environment for model building into low-resolution electron-density maps. Acta Crystallogr D Struct Biol 74, 519–530, doi:10.1107/S2059798318002425 (2018).

41 Emsley, P. & Cowtan, K. Coot: model-building tools for molecular graphics. Acta Crystallogr D Biol Crystallogr 60, 2126–2132, doi:10.1107/S0907444904019158 (2004).

42 Adams, P. D. et al. Recent developments in the PHENIX software for automated crystallographic structure determination. J Synchrotron Radiat 11, 53–55, doi:10.1107/s0909049503024130 (2004).

43 Davis, I. W., Murray, L. W., Richardson, J. S. & Richardson, D. C. MOLPROBITY: structure validation and all-atom contact analysis for nucleic acids and their complexes. Nucleic Acids Res 32, W615–619, doi:10.1093/nar/gkh398 (2004).

44 Barad, B. A. et al. EMRinger: side chain-directed model and map validation for 3D cryo-electron microscopy. Nat Methods 12, 943–946, doi:10.1038/nmeth.3541 (2015).

45 Goddard, T. D. et al. UCSF ChimeraX: Meeting modern challenges in visualization and analysis. Protein Sci 27, 14–25, doi:10.1002/pro.3235 (2018).

46 Chan, J. F. et al. Simulation of the clinical and pathological manifestations of Coronavirus Disease 2019 (COVID-19) in golden Syrian hamster model: implications for disease pathogenesis and transmissibility. Clin Infect Dis, doi:10.1093/cid/ciaa325 (2020).

47 Chan, J. F. et al. Improved Molecular Diagnosis of COVID-19 by the Novel, Highly Sensitive and Specific COVID-19-RdRp/Hel Real-Time Reverse Transcription-PCR Assay Validated In Vitro and with Clinical Specimens. J Clin Microbiol 58, doi:10.1128/JCM.00310-20 (2020).

